# Light-induced structural adaptation of the bundle-shaped phycobilisome from thylakoid-lacking cyanobacterium *Gloeobacter violaceus*

**DOI:** 10.1101/2025.05.12.653420

**Authors:** Jianfei Ma, Xin You, Shan Sun, Sen-Fang Sui

## Abstract

*Gloeobacter* diverged from other lineages early in cyanobacterial evolution, preferentially growing under low light intensity conditions. Among cyanobacteria, *G. violaceus* exhibits unique features, including lack of a thylakoid membrane and bundle-shaped antenna phycobilisomes (PBSs), densely packed and well-organized on the plasma membrane. However, without high-resolution structures, it has remained unclear how *G. violaceus* PBSs assemble into a bundle-shaped configuration. Here we solved the cryo-EM structures of PBSs from *G. violaceus* cells cultured under low (Sr-PBS) or moderate (Lr-PBS) light intensity. These structures revealed two unique linker proteins, L_RC_^91kDa^ and L_RC_^81kDa^, that play a key role in the PBS architecture. Analysis of the bilin arrangement indicated that the bundle-shaped structure allows efficient energy transfer among rods. Moreover, comparison between Lr-PBS and Sr-PBS uncovered a distinct mode of adaption to increased light intensity wherein the ApcA_2_-ApcB_3_-ApcD layer can be blocked from binding to the core by altering structural elements exclusively found in the *G. violaceus* L_CM_. This study illustrates previously unrecognized mechanisms of assembly and adaptation to varying light intensity in the bundle-shaped PBS of *G. violaceus*.

## Introduction

*Gloeobacter* clusters on the most basal branch of the cyanobacterial phylogenetic tree and phylogenetic analyses based on 16S ribosomal RNA (rRNA)^1^, some housekeeping genes^2–4^ and/or thylakoid morphology and architecture^5, 6^ suggest this genus may be closest to the most ancient cyanobacteria, supporting its value for study of the evolutionary history of oxygenic photosynthesis. In particular, the rod-shaped unicellular cyanobacterium, *Gloeobacter violaceus* PCC 7421 is the most widely studied *Gloeobacter* species. *G. violaceus* was first isolated from the surface of limestone in Switzerland^7^, and subsequently detected in environmental conditions considered marginal for other cyanobacteria^8, 9^. Consistent with its basal position in the phylogenetic tree and distinct niche, *G. violaceus* exhibits several unique traits, notably lacking a multiple, stacked thylakoid membrane structure^10, 11^. The photosynthesis and respiratory apparatus is therefore required to operate within the single layer of the cytoplasmic membrane^12^, which consequently limits its metabolic and growth rates^13, 14^. Additionally, compared to crown cyanobacteria, the *psbQ*, *psbY*, *psbZ* and *psb27* genes in photosystem II (PSII) are absent, while three genes encoding extrinsic proteins (PsbO, PsbU and PsbV) are poorly conserved^11^. Moreover, photosystem I (PSI) contains a reduced number of subunits in *G. violaceus*^15^. The relatively low concentrations of carotenoids in this species, equivalent to about 10% of total carotenoids in *Synechocystis* ^16, 17^, results in limited protection for the photosynthetic apparatus from high light intensities, which might also contribute to its slow growth under strong light. Previous work has demonstrated that development of *G. violaceus* cells is adversely affected by high light intensity, with a maximum tolerable intensity of approximately 100 fc (∼ 40 μmol of photons m^−2^ s^−1^)^12^.

Similar to other cyanobacteria, the major light harvesting complex in *G. violaceus* is phycobilisome (PBS), which exhibits a distinctive bundle shape that is highly organized on the cytoplasmic membrane, rather than the hemidiscoidal conformation common in other cyanobacteria^18^. In general, PBS consists of phycobiliproteins (PBPs), including phycocyanin (PC), phycoerythrin (PE) and allophycocyanin (APC), along with linker proteins, and a wide diversity of open-chain tetrapyrrole chromophores that covalently bind to PBPs and some linker proteins to mediate light absorption and energy transfer^19–21^. Structural analysis has shown that a typical PBS contains a central core comprising an APC surrounded by several peripheral PC and PE rods^22–26^, with linker proteins that form a scaffold to ensure the ordered and accurate assembly of the entire PBS^27^. Genomic studies have shown that *G. violaceus* contains seven types of PBPs (ApcA, ApcB, CpcA, CpcB, CpeA, CpeB and ApcD), lacking only ApcF^11, 14, 28^. However, despite the presence of *apcD* in the *G. violaceus* genome, no ApcD protein has yet been identified among its purified phycobiliproteins^29^. It therefore remains unclear whether ApcD itself is absent from the *G. violaceus* PBS in *vivo* or if it is lost during the purification/isolation process. Among the 11 linker proteins detected in *G. violaceus*, glr1262 (81.4 kDa) and glr2806 (91.9 kDa) are larger than other linker proteins except the core-membrane linker (L_CM_ or ApcE), and have not yet been found in the PBS of any other species. Both of these linkers contain three tandem repeats of the Pfam00427 domain, while other linker proteins, except L_CM_, contain only a single Pfam00427 domain or lack it altogether. The location and function of glr1262 and glr2806 have long remained controversial^28, 30, 31^ until recent work identified them as rod-core linkers. Mutant analysis together with biochemical assays and electron microscopy conducted by Wang and coworkers suggest that glr1262 is responsible for the attachment of bundled rods to the core, while glr2806 connects additional rod hexamers to the core^32^. However, the mechanism through which these two linker proteins attach rods to the core to facilitate formation of the distinct bundle-shaped PBS is still unclear.

To address the above questions, in the present study, we cultured *G. violaceus* 7421 under low and moderate light conditions, then employed cryo-electron microscopy (cryo-EM) to determine the PBS structure at overall resolutions of 3.37 Å and 2.8 Å, respectively uncovering long rods that contain an ApcA2-ApcB3-ApcD layer and short rods that lack this layer. These structures depict the detailed organization between linker proteins and surrounding PBPs, providing a mechanistic understanding of the process leading to formation of the bundle-shaped PBS, as well as structural regulation of light-induced binding to the ApcA2-ApcB3-ApcD layer. These high resolution structures also reveal a distinct chromophore distribution corresponding to the distinct bundle shape and suggest possible energy transfer pathways in this primitive cyanobacterial PBS.

## Results and Discussion

### Two bundle-shaped PBS structures from *G. violaceus*

To observe structural adaptations of the PBS to different environments, *G. violaceus* 7421 cells were cultured under low and moderate light intensities (Abbreviated as LL and ML) (∼5 μmol and ∼35 μmol photons m^−2^ s^−1^, respectively) with a 16 h:8 h light– dark cycle at 20°C to mimic its natural growth conditions. Under these conditions the purified PBS exhibited purple and blue colors, respectively, which corresponded well with their absorption spectra (Supplementary Fig. 1a). Three absorbance peaks (∼500 nm, 560 nm and 630 nm) were detected for the purple PBS, while blue PBS showed only one absorbance peak at 630 nm, indicating that PE was absent from the *G. violaceus* PBS following growth under ML intensity (Supplementary Fig. 1b). We subsequently applied single-particle cryo-EM analysis to resolve the structures of these PBSs obtained from LL and ML *G. violaceus* cells, at overall resolutions of 3.37 Å and 2.8 Å, respectively. The most prominent difference we observed between the two structures was that PBS from ML *G. violaceus* cells contained distinctly shorter rods (hereafter, short-rod PBS, Sr-PBS) compared to PBS isolated from LL *G. violaceus* cells (long-rod PBS, Lr-PBS). The Lr-PBS (14.0 MDa) had a height of ∼530 Å, while the Sr-PBS (7.0 MDa) had a height of ∼406 Å, and both exhibited a width of ∼438 Å (Fig. 1a and b). Applying individual local masks to reconstruct the Sr-PBS improved the resolutions of rods a-d (Ra-Rd) to 2.6 - 3.5 Å (Supplementary Fig. 2). Given that the peripheral densities of the rods were indistinct, atomic models were only constructed for residues and bilins in the high-resolution regions, including the APC core, the 1^st^ and the 2^nd^ hexamers of the rods in Sr-PBS, and the APC core of Lr-PBS (Supplementary table. 1).

**Figure 1.**
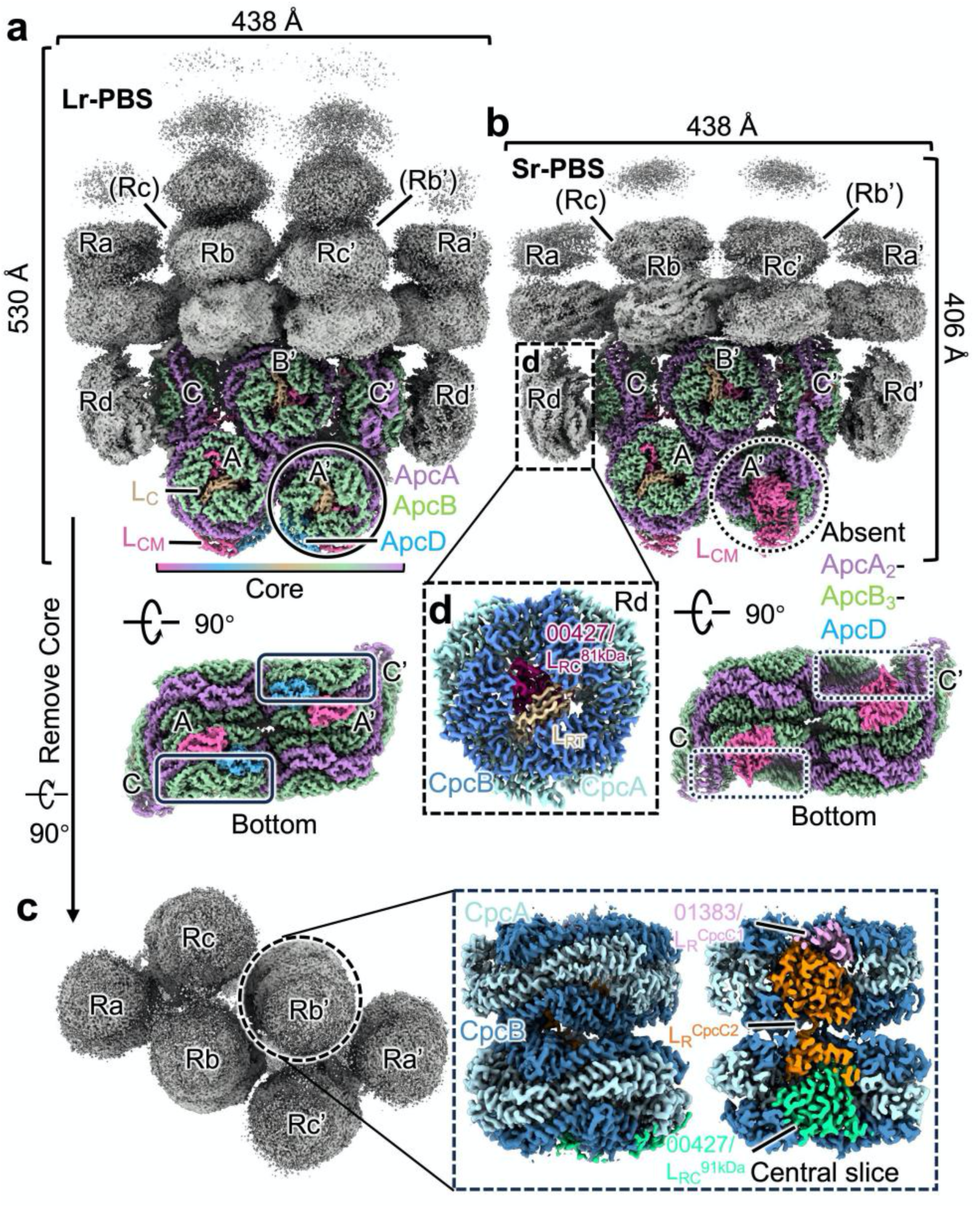
Overall structures of PBS from *G. violaceus* 7421. **a-b**, Cryo-EM maps of two types of assemblies of PBS (Lr-PBS and Sr-PBS) from face and bottom views. The rods are colored gray. PBS core is presented in different colors. The solid and dash black circles are represented whether ApcA_2_-ApcB_3_-ApcD layer is bond with core or not in Lr-PBS and Sr-PBS. **c**, Top view of rods. Enlarged view of the boxed area on the right side of **c** shows the cryo-EM map of Rod b after local refinement. The L_R_^CpcC1^, L_R_^CpcC2^ and L_RC_^91kDa^ are colored as pink, orange and limegreen. **d**, Enlarged view of the boxed area of **b** shows the cryo-EM map of Rod d after local refinement. The L_RT_ and L_RC_^81kDa^ are colored as wheat and warmpink.

The APC core of both Lr-PBS and Sr-PBS comprised 5 cylinders, including a top cylinder (B) stacked above two basal cylinders (A and A’), and two half-cylinders (C and C’) positioned vertically on the shoulders of A and A’ (Fig. 1a and b). This conformation was in agreement with the pentacylindrical PBS core reported for *Anabaena* 7120 ^24^. Interestingly, compared to Lr-PBS, the outermost APC trimers of cylinders A and A’ (ApcA_2_-ApcB_3_-ApcD layer) were absent in the Sr-PBS (Fig. 1a and b). As this difference was associated with a change in ambient light intensity from ∼5 μmol to ∼35 μmol photons m^−2^ s^−1^, we hypothesized that the PBS structure from *G. violaceus* might undergo adaptive changes in response to varying light intensity.

The eight peripheral rods (Ra/a’ - Rd/d’) were subsequently classified into two types based on their positions relative to the APC core. Six rods (Ra/a’- Rc/c’) were designated as Type I based on their upright placement stacked above the core, resulting in a so-called “bundle-shaped” conformation (Fig. 1 a-c). Although medium-resolution maps showed as many as seven hexameric layers of type I rods, the layers distal to the core were too dynamic, and therefore we could only estimate the number of hexameric layers per rod in subsequent data processing. Only five and three hexameric layers could be clearly observed in the final high-resolution reconstructions of Lr-PBS and Sr-PBS rods, respectively (Fig. 1 and Supplementary Fig. 1a). The extended hexamers of Lr-PBS were likely PEs, which was previously reported based on negative staining in a △PE-PBS mutant^32^.

In contrast, Type II rods (Rd/Rd’) were positioned laterally along the side of the core C/C’, forming a platform to support Ra/a’ (Fig. 1a and b). These Type II rods contained only one PC hexamer, potentially due to the location of the rod-terminal linker protein (L_RT_) within the cavity of the PC hexamer, which can reportedly inhibit rod elongation^33^ (Fig. 1d). In addition, low-pass filtration resulted in blurred densities, symmetrically distributed close to the APC cylinders A(A’) and rod Rd (Rd’) (Supplementary Fig. 1a). Although it is challenging to identify the corresponding protein components due to limitations imposed by resolution, it is plausible that this electron dense region may represent a lateral rod (defined as Re/e’), and considering its position adjacent to rod Rd (Rd’), may be connected by a shared rod-core linker protein, L_RC_^81kDa 32^ (Supplementary Fig. 1). In addition to the core and rods, we determined the structures of several linker proteins, including two rod linker proteins (L_R_^Cpc1^ and L_R_^Cpc2^), one rod-terminal linker L_RT_, two rod-core linker proteins (L_RC_^91kDa^ and L_RC_^81kDa^) with molecular masses of 91.9 kDa and 81.4 kDa, respectively, the core linker protein (L_C_), and the core–membrane linker protein (L_CM_) (Supplementary Fig. 3).

Assembly of the APC core depends on the L_CM_ structure, which has multiple Pfam00427 domains for anchoring APC cylinders, thereby forming bicylindrical, tricylindrical, and pentacylindrial-typed APC cores^34, 35^. The L_CM_ from *G. violaceus* contains 4 Pfam00427 domains, which is the same as that reported in another ancient cyanobacterium *Anthocerotibacter panamensis* (*A. panamensis*) that diverged from *Gloeobacter* spp. around 1.4 billion years ago^14^, and the crown cyanobacterium *Anabaena* 7120^24^. Superpositioning of the L_CM_ protein structures from *G. violaceus*, *A. panamensis*, *Anabaena* 7120, *Synechococcus* 7002, and *P. purpureum* indicated that Pfam00427 domains 1-3 (00427-1 to 00427-3) aligned well across these cyanobacterial species, while the Pfam00427 domain 4 (00427-4) showed relatively high similarity among *G. violaceus*, *A. panamensis,* and *Anabaena* 7120 (Supplementary Fig. 4). This structural conservation suggests that the L_CM_ maintained a stable conformation throughout its evolution, and the assembly mode of the APC core was present in the common ancestor of these species.

### Bundle-shaped rod assembly mediated by L_RC_^91kDa^ and L_RC_^81kDa^

As *G. violaceus* strikingly lacks a thylakoid membrane characteristic of crown cyanobacteria, PBSs are necessarily confined to the plasma membrane. The bundled shape could thus facilitate dense packing of PBSs into the limited membrane area. Previous studies suggest that L_RC_ could mediate interactions between the basal hexamers in rods and the core, consequently determining the direction in which each rod extends^36^. However, *G. violaceus*, lacks the canonical the L_RC_ gene, *CpcG,* but instead harbors, *glr1262* and *glr2806*, encoding L_RC_^91kDa^ and L_RC_^81kDa^, respectively^22, 37–39^. As neither L_RC_^91kDa^ nor L_RC_^81kDa^ have been found in crown cyanobacteria or red algae, their structures and functions remain unclear. L_RC_^91kDa^ contains three Pfam00427 domains (Pfam00427-1 to Pfam00427-3) tethered in a triangular pattern by three loop regions (loop-1 to loop-3), positioned above the top core cylinders, B/B’ and C (Fig. 2a and b). These loop regions wrap around the surfaces of the cores and interact with grooves in the α-subunits via four short helices, primarily through conserved hydrophobic residues. These interactions are similar to those observed between L_RC_ proteins and the PBS core in crown cyanobacteria and red algae^22–24, 26^, implying that this binding pattern may have originated in a common ancestor (Fig. 2b and c). Although CpcN, from the thylakoid-free cyanobacterium, *A. panamensis,* contains five Pfam00427 domains, each Pfam00427 domain is anchored by L_RT_ to inhibit rod elongation^40^. This interaction differs from that of L_RC_^91kDa^, which we found was attached to the Pfam01383 domain of L_R_^Cpc2^. Furthermore, L_RC_^91kDa^ shares low sequence identity with CpcN (44.1%) in in NCBI BLASTP results, nor was close to CpcN in phylogenetic analysis (Fig. 3a), suggesting that L_RC_^91kDa^ is not likely a homolog of the rod-core linker, CpcN. Thus, speculated that the presence of L_RC_^91kDa^ exclusively in the *G. violaceus* lineage might contribute to its unique bundle-shaped assembly.

**Figure 2.**
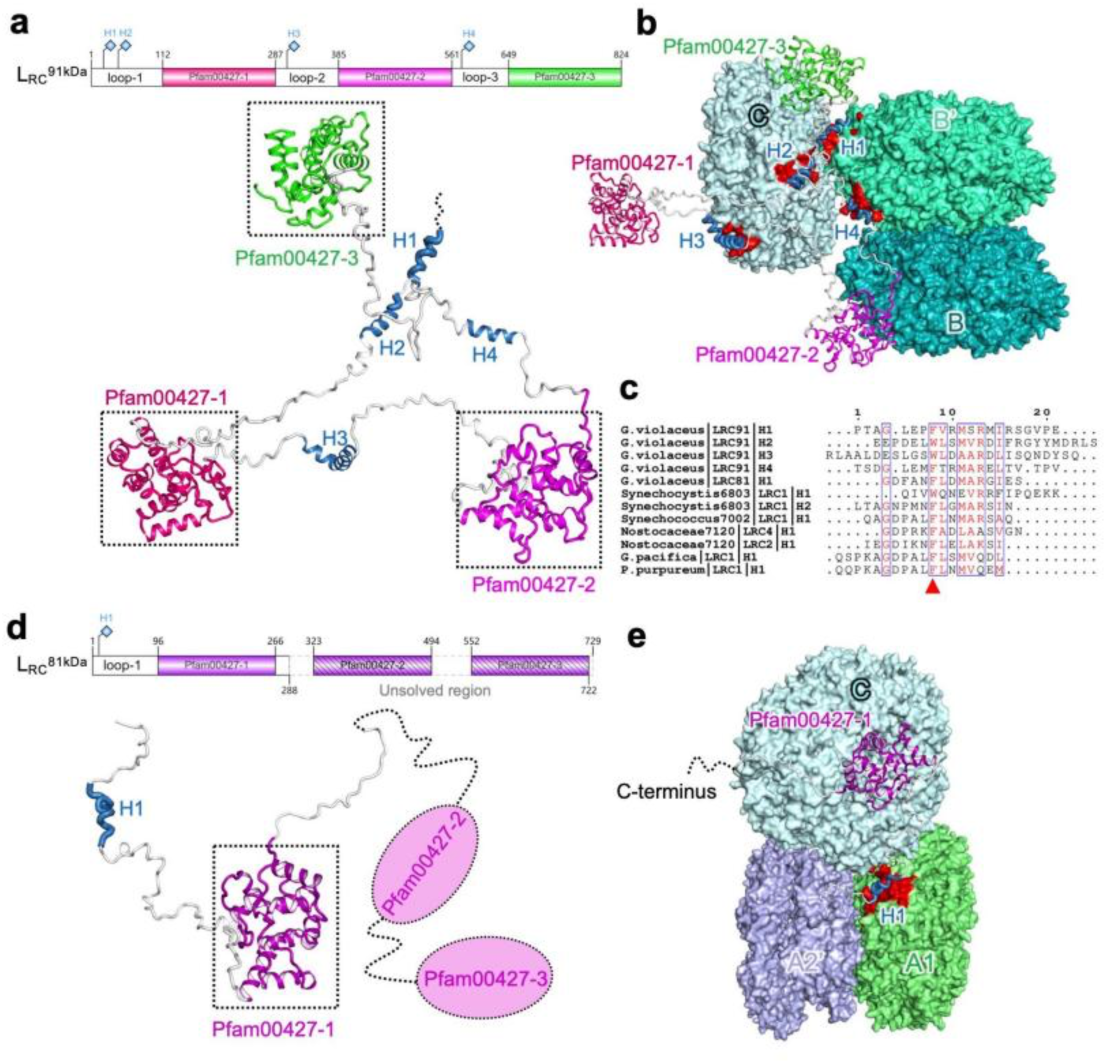
Rod-core linker proteins, L_RC_^91kDa^ and L_RC_^81kDa^. **a**, Top, diagram of the structural elements of L_RC_^91kDa^. Bottom, the structure of L_RC_^91kDa^ is represented as cartoon. Three Pfam00427 domains are presented in different colors. Four helices are highlighted in skyblue. **b**, Interactions between L_RC_^91kDa^ and core. The grooves on the α-subunits that contact the helices are shown in red. **c**, Sequence alignment of the helices of L_RC_^91kDa^ with L_RC_ from *Synechocystis* sp. PCC 6803, *Synechococcus* sp. strain PCC 7002, *Nostocaceae* PCC 7120, *G.pacifica* and *P.purpureum*. The conserved Phenylalanine or Tryptophan is marked as red triangle. **d**, Top, diagram of the structural elements of L_RC_^81kDa^. Bottom, the structure of L_RC_^91kDa^ is represented as cartoon. The Pfam00427-1 domain we solved is presented in megentas. The Pfam00427-2 and -3 are is presented in dash ellipse. The helix is highlighted in skyblue. **e**, Interactions between L_RC_^81kDa^ and core. The groove on the α-subunit that contact the helix is shown in red.

**Figure 3.**
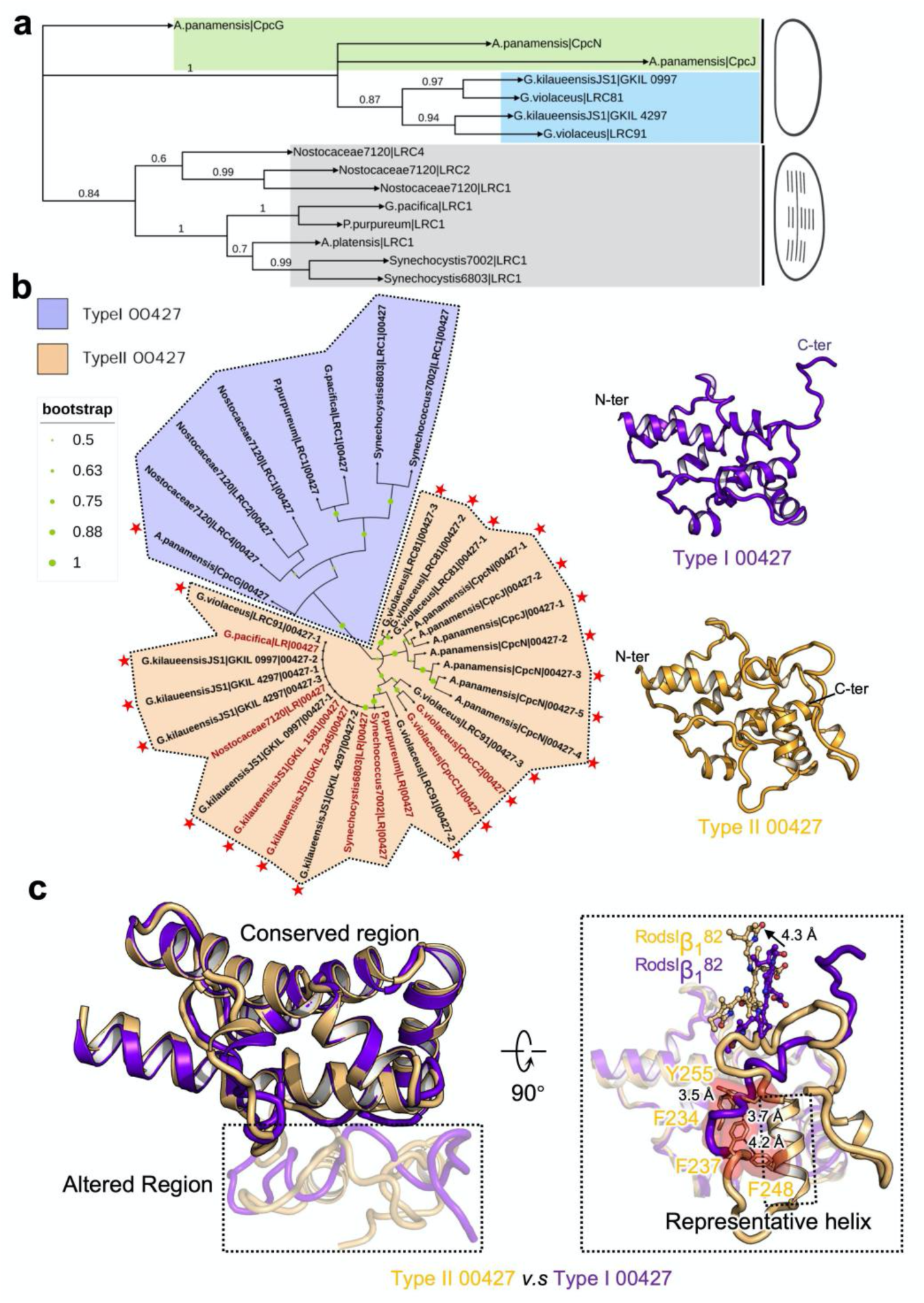
Phylogenetic relationship of L_RC_ and classifications of Pfam00427. a, The maximum likelihood phylogenetic trees constructed using the full-length protein sequences of L_RC_ from thylakoid-lacking cyanobacterium *G. violaceus* 7421, *G. kilaueensis* JS1, *A. panamensis*, and crown cyanobacteria *Nostocaceae* PCC 7120, *Synechocystis* sp. PCC 6803, *Synechococcus* sp. strain PCC 7002, *A. platensis* and red algae *G.pacifica* and *P.purpureum*. Bootstrap values are presented on the tree nodes, and the values higher than 0.6 are shown. b, Left, phylogenetic trees constructed using the separated Pfam00427 domains, Pfam00427 domains of L_R_ and L_RC_ are labeled in firebrick and black, respectively. The Pfam00427 domains are classified to two types: Type I and II 00427 domains are shown on the purple and light orange background, respectively. The Pfam00427 domains belong to thylakoid-free cyanobactera are marked with red star. Right, the structures of type I and II 00427 domains are shown as cartoon. c, Structural alignment of the type I and II 00427 domains. Region of structural change is shown with 70% transparency. Enlarged view of the boxed area of c shows the altered structures including key aromatic residues, representative helix of type II 00427 domain, as well as the shift of bilins.

L_RC_^81kDa^ also comprises three Pfam00427 domains, which we found connected the lateral rods (Rd/d’) to the core via interactions between the α-helix and α-subunit (Fig. 1, Fig. 2c - e). These interactions resulted in the lateral Rd/d’ forming a platform to support the Ra/a’-Rc/c’ in the bundle-shaped conformation (Fig. 1). Unfortunately, only the 00427-1 domain was clearly resolved in our structure, possibly due to the structural dynamics of the 00427-2 and 00427-3 domains. L_RC_^81kDa^ bound to the lateral side of the APC cylinder and bound to L_RT_ to block extension of Rd/d’, similar to CpcJ protein in *A. panamensis* ^40^. However, the low sequence identity (34.9%) and low relatedness in our phylogeny suggested that they may have diverged independently during the evolution of PBS (Fig. 3a).

L_R_ proteins are known to play important roles in rod extension^22, 39^. L_R_^Cpc2^ contained an N-terminal Pfam01383 domain (M1-E75) and a C-terminal Pfam00427 domain (D101-V274) connected by a hinge (A76-S100) structure (Supplementary Fig. 5a). The Pfam00427 domain was located in cavities formed by the 2^nd^ hexamers in rods Ra/a’- c/c’, whereas the Pfam01383 domain extended into basal hexamers and directly contacted the Pfam00427 domain of L_RC_^91KDa^. Structural alignments of the Pfam01383 domain showed that although L_R_^Cpc2^ exhibited structural features similar to those of the L_R_ in crown cyanobacteria and red algae, its Pfam00427 domain was shifted ∼24 Å and rotated ∼120° (Supplementary Fig. 5b). This difference in rod orientation was likely due to extension of the L_R_ and L_R_^Cpc2^ hinges in opposite directions, resulting in a correspondingly deflected position of the linked Pfam00427 domain. This shift in Pfam00427 domain orientation could guide the direction in which associated hexamers are extended, circumventing excessive protrusion of Ra/a’-c/c’ (Supplementary Fig. 5c). Thus, L_R_^Cpc2^ might also serve as a contributing factor in the formation of bundle-shaped rods.

### Classification of Pfam00427 domains

Phylogenetic analysis showed that L_RC_s in cyanobacteria and algae evolved independently between those with or without thylakoid membranes (Fig. 3a). For example, L_RC_^91kDa^ and L_RC_^81kDa^ shared a close relationship with GKIL_4297 and GKIL_0997 from *G. kilaueensis* JS1, but appeared distantly related to CpcG, CpcN and CpcJ from *A. panamensis*, suggesting that the PBS in *G. kilaueensis* JS1 likely exhibits a bundle shape. Interestingly, the Pfam00427 domains of various L_RC_s and L_R_s from thylakoid-free cyanobacteria, crown cyanobacteria and red algae could be divided into two types based on their sequence similarity. We found type I Pfam00427 only in L_RC_s that contained a single Pfam00427 domain, whereas type II Pfam00427 was present in both L_R_s and L_RC_s with multiple Pfam00427 domains (Fig. 3b).

Superpositioning of the type I and type II Pfam00427 structures showed that they both harbored a conserved region that could bind to the Pfam01383 domain, as well as an altered region at the C-terminus (Fig. 3c). Different from the C-terminal loop in type I Pfam00427, the C-terminus of type II Pfam00427 could fold into a typical helix, wherein four aromatic residues (F234, F237, F248 and Y255) appear stacked in a sandwich orientation to stabilize its conformation. These residues are characteristically conserved in type II Pfam00427, but replaced with polar residues (e.g., Q234, R237 and Q255) in type I Pfam00427. Among these residues, F237, located at the start of the altered region, was strictly conserved among type II Pfam00427 domains from different species (Fig. 3c, Supplementary Fig. 6a and b). We thus speculated that F237 could potentially play a key role in the type II Pfam00427 structure. In contrast, the presence of R237 in place of F237 could potentially disrupt such sandwich-stacking in type I Pfam00427 domains, leading to a disordered loop structure. Correspondingly, the pigment, ^RodsI^β_1_^82^, in type II Pfam00427 exhibits a 4.3 Å shift relative to that in type I Pfam00427 to avoid steric clashes (Fig. 3c). Moreover, the rod hexamer bound to type I Pfam00427 was deflected by ∼20° relative to rod hexamers bound to type II Pfam00427, consequently altering the position and orientation of the corresponding pigments (Supplementary Fig. 6c), which could affect the spectroscopic properties of the pigments.

Phylogenetically, type I Pfam00427 of CpcG from *A. panamensis* clustered together with that from crown cyanobacteria and red algae (Fig. 3b). Given that *A. panamensis* is a relatively ancient cyanobacterium and the *cpcG* gene is absent in *Gloeobacteria* lineages, it was reasonable to speculate that L_RC_s containing only a single Pfam00427 domain might have evolved from the CpcG in *A. panamensis*. As type II Pfam00427 of L_R_s showed a scattered distribution among L_RC_s that contain multiple Pfam00427 domains, with no obvious genetic relationship among them, the evolutionary trajectory of L_R_s in crown cyanobacteria and red algae currently remains uncertai.. For instance, it is possible that these type II domains represent evolutionary relics retained from an ancestral L_R_ of *G. violaceus*, or alternatively, gradually diverged from L_RC_s containing a type II Pfam00427.

### Preferential interactions of β^82^s with linker proteins in *G. violaceus*

Previous studies have shown that sunlight absorbed by peripheral rods of PBS can be funneled to the core via bilins in remarkably high efficiency^41^. In this study, the PCB was the only bilin identified in Lr-PBS or Sr-PBS structures, as PEB was located in the blurred (i.e., unsolved) structures of distal PE hexamers. A total of 336 PCBs were identified in Sr-PBS, 252 of which were located in rods and 84 were identified in the core, while Sr-PBS contained only 96 total PCBs (Supplementary Table. 2). For each hexamer unit, energy absorbed by the exterior bilins has been shown to be subsequently transferred to the three β82 bilins located adjacent to the inner cavity^42^.

Previous structural observations^23^ and quantum chemical analysis^43^ suggest that these β82s undergo preferential interactions with linker proteins, resulting in different energy states. Based on this prior knowledge, we explored β82-linker interactions in Rbs. Examination of Rb2I showed that β82 PCBs (designated as ^Rb2I^β_1_^82^, ^Rb2I^β_2_^82^, and ^Rb2I^β_3_^82^) were distributed around L_R_^Cpc2^ (Fig. 4a and b), with three aromatic residues of L_R_^Cpc2^ (F224, F235 and F239) stacked in a T-shaped conformation adjacent to the pyrrole B ring of ^Rb2I^β_2_^82^, forming a π electron platform to stabilize the ring moiety^44^. In contrast, no aromatic residues were located close to the ^Rb2I^β_1_^82^, and only the Y118 of L_R_^Cpc2^ forming a T-shaped structure to mediate π-π interaction with ^Rb1I^β_3_^82^ at a distance of 4.5 Å (Fig. 4c-e). Consistent with the previous studies^23, 24, 43^, this organization suggests that ^Rb2I^β_2_^82^ like exhibits the lowest energy level compared to ^Rb2I^β_1_^82^ and ^Rb2I^β_3_^82^. Similar features were also observed in the core-proximal PC hexamers (Fig. 4f), wherein four aromatic residues (F510, Y512,F521 and F528) of the L_RC_^91kDa^ 00427-2 domain formed a π electron platform positioned closely (4.3 Å) to ^Rb1I^β_1_^82^, while only Y404 and F474 were respectively located around ^Rb1I^β_2_^82^ and ^Rb1I^β_3_^82^ (Fig. 4g - i).

**Figure 4.**
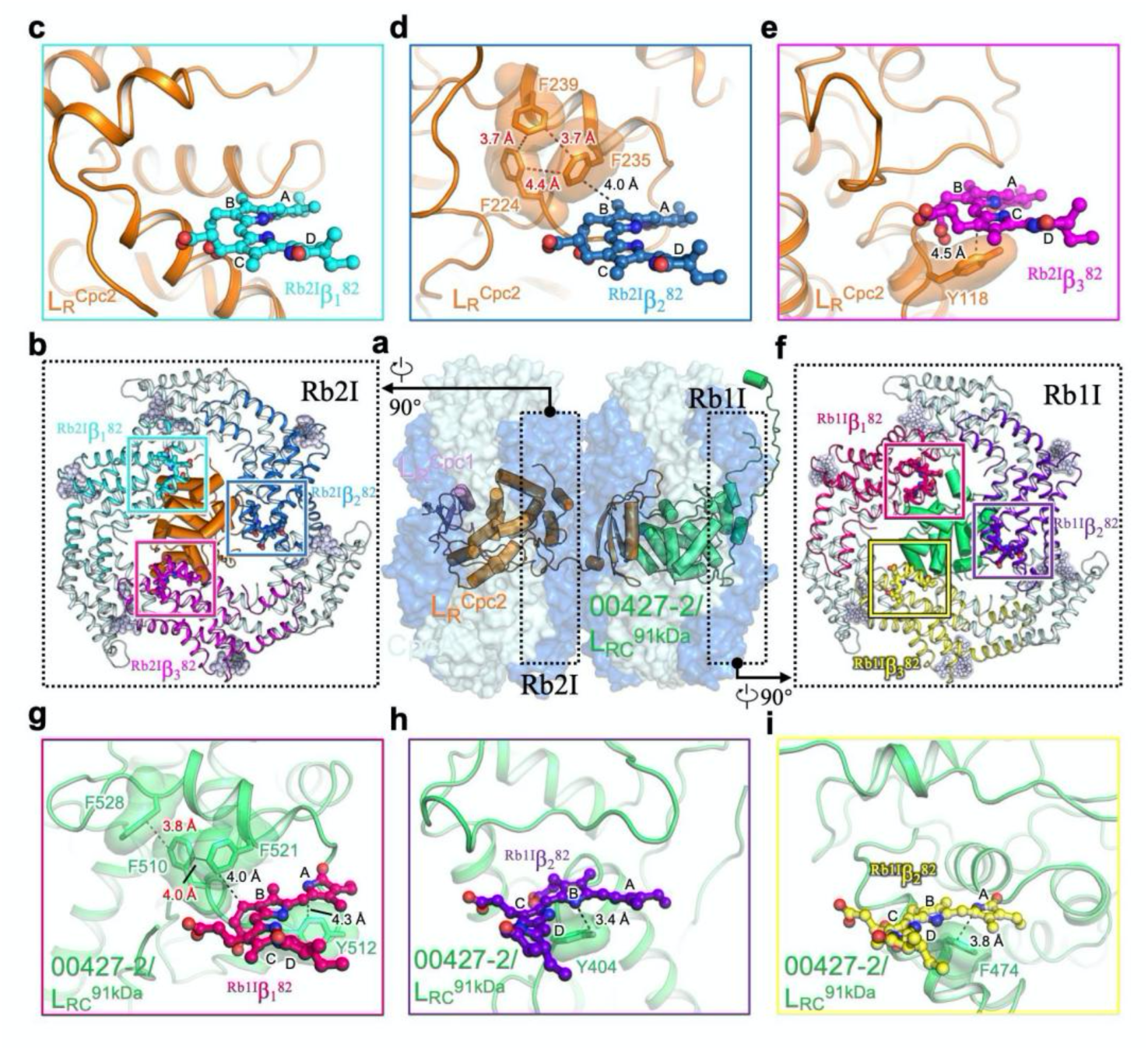
Interactions of the linker proteins L_R_^CpcC2^ and L_RC_^91kDa^ with chromophores in the rod Rb. **a**, Overall structure of the rod Rb with the hexamers shown in surface representation and the linker proteins shown in cartoon representation. **b**, Structure of the layer Rb2I. Proteins and bilins are shown in cartoon and sticks representations, respectively. Three β subunits are coloured differently and the β82 PCBs are boxed and analysed in **c-e**. **c**, No interactions between the L_R_^CpcC2^ and the bilin ^Rb2I^β_1_^82^. **d**, The interactions between F224, F235 and F239 and the bilin ^Rb2I^β_2_^82^. **e**, The interactions between Y118 and ^Rb2I^β_3_^82^. **f**, Structure of the layer Rb1I. Proteins and bilins are shown in cartoon and sticks representations, respectively. Three β subunits are coloured differently and the β82 PCBs are boxed and analysed in **g-i**. **g**, The interactions between F510, F512, F521 and F528 and the bilin ^Rb1I^β_1_^82^. **h**, The interactions between Y404 and the bilin ^Rb1I^β_2_^82^. **i**, The interactions between F474 and the bilin ^Rb1I^β_3_^82^.

Examination of the lateral Rd1 showed that ^Rd1I^β_3_^82^ might also formed π-π interactions with F221, Y223 and F232 of the L_RC_^81kDa^ 00427-1domain, whereas ^Rd1I^β_1_^82^ and ^Rd1I^β_2_^82^ each respectively formed interactions with only a single Pfam00427-1 domain residue (Supplementary Fig. 7a and b). Sequence alignment showed that these aromatic residues were well conserved among type II Pfam00427 domains of L_R_s and L_RC_s, suggesting that regulation of the β82 bilin energy state via aromatic residues likely originated in ancient cyanobacteria (Supplementary Fig. 6a). Notably, these β^82^ PCBs were located closest to the core in Ra1I – Rd1I, further suggesting that they could act as an energy sink to coordinate energy migration from the rods and core (Supplementary Fig. 8a and b). A similar situation was previously identified in rods of PCBs in the red algae, *P. purpureum* and *G. pacifica,* further supporting the high evolutionary conservation of this mechanism^22, 23, 43^.

### Binding specificity of the ApcA_2_-B_3_-ApcD layer in *G. violaceus*

ApcD is important for energy transfer from PBS to PSII and PSI^45^, and stably binds to the bottom cylinder of the PBS core in crown cyanobacteria and red algae^22–24^. In contrast, we observed that the bundle-shaped PBS of *G. violaceus* associated with the ApcA_2_-ApcB_3_-ApcD trimer under low intensity light, but disassociated from the Apc trimer under moderate light intensity (Fig. 1 and Supplementary Fig. 1). The ApcA_2_-ApcB_3_-ApcD trimers bound to the A and A’ core cylinders via the first Pfam00427 domain of L_CM_/L_CM_’ and the L_C_/L_C_’ core linker. Sequence alignment of L_CM_ from various cyanobacteria and red algae revealed two extra loops (termed as loop1 and loop2) that were only present in the first Pfam00427 domain of the *G. violaceus* L_CM_ (Supplementary Fig. 4 and 9). Structural analysis indicated that loop1 and loop2 exhibited obviously different conformations between Lr-PBS and Sr-PBS (Fig. 5a). In particular, R283 and K285 in loop1 of Lr-PBS_LCM_ swings toward the ApcA subunit, forming several electrostatic and polar interactions with residues D105, N110 and E114 of ApcA (Fig. 5b and c). However, as loop1 is rotated upwards (consider including a degree of change) in Sr-PBS_LCM_, these interactions with ApcA are abolished, potentially leading to dissociation from the ApcA2-ApcB3-ApcD trimer (Fig. 5b). In addition, Pfam00427 domain loop2 of Lr-PBS_LCM_ also rotates around to form several polar interactions with the L_C_. However, loop2 of Sr-PBS_LCM_ occupies the C-terminal position of the L_C_, resulting steric hindrance that obstructs binding with the C-terminus of L_C_ (Fig. 5d). As L_C_ is crucial for assembly of the APC trimer^38, 46^, it is likely that such a conformational change in Sr-PBS_LCM_ loop2 could impede assembly of the ApcA_2_-ApcB_3_-ApcD layer. Collectively, these results suggest that loop1 and loop2 of the first Pfam00427 domain in L_CM_ may function together with L_C_ to act as a “switch” in *G. violaceus* to regulate binding with the ApcA_2_-ApcB_3_-ApcD trimer in response to changes in light intensity. Notably, ApcD is absent in the *A. panamensis Gloeobacterial* lineage, implying that unstable binding between ApcD and the PBS core may represent an ancestral state^40^.

**Figure 5.**
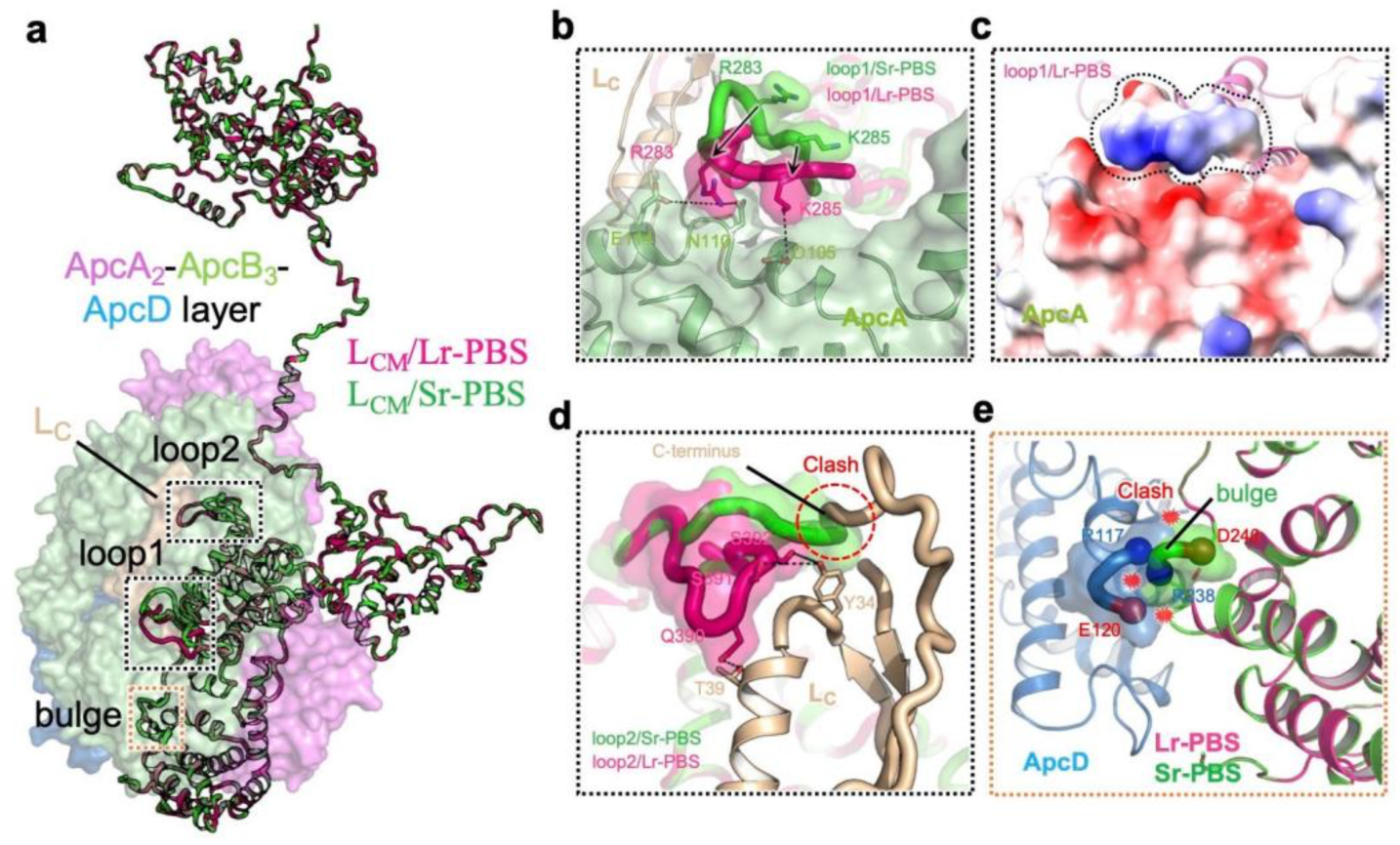
Structural differences between L_CM_/Lr-PBS and L_CM_/Sr-PBS. **a**, Structural alignment of L_CM_/Lr-PBS and L_CM_/Sr-PBS. L_CM_/Lr-PBS and L_CM_/Sr-PBS are presented in hotpink and green color. ApcA2-ApcB3-ApcD layer attached with L_CM_/Lr-PBS is shown as surface view. The altered structures (loop1, loop2 and bulge) are highlighted in dash rectangles. **b**, Enlarged view of the boxed area of **a** shows the structural change of loop1 reduces the interactions between loop1 and ApcA. **c**, Electrostatic interaction between loop1/Lr-PBS and ApcA. **d**, Enlarged view of the structural change of loop2 shows the absent of the interactions between loop2/Sr-PBS and L_C_. loop2/Sr-PBS could form a steric hindrance to prevent L_C_ binding with the core. **e**, Three successive residues (R238-S239-D240) of α^LCM^ could form a “bulge” that would be occupied the position of the ApcD.

Moreover, we noted that the α domain of L_CM_ (α^LCM^) contains three successive residues (R238-S239-D240) in *G. violaceus* that are absent in the α^LCM^ domain of crown cyanobacteria and red algae (Supplementary Fig. 9). In the α^LCM^ domain of Lr-PBS_LCM_, the EM density representing these residues disappeared, potentially creating a space that could allow ApcD binding to the PBS core. Conversely, in the α^LCM^ of Sr-PBS_LCM_, the density was clearly solved, forming a “bulge” at the same position occupied by ApcD in Lr-PBS_LCM_, consequently preventing ApcD binding with the PBS core (Fig. 5e). Although we cannot completely exclude the possibility of trimer loss during purification, we observed the clearly solved densities representing loop1, loop2 and the bulge in L_CM_ that could all obviously participate in ApcA_2_-ApcB_3_-ApcD trimer binding to the PBS core.

### Bilin arrangement and possible energy transfer pathways

The hemidiscoidal-shaped PBS from crown cyanobacteria exhibits a dispersed arrangement of rods, resulting in minimal energy transfer among rods^26^. However, in *G. violaceus* the dense packing of rods in the bundle-shaped PBS result in a distance of 30 Å or less between the outermost PCBs of adjacent rods, which could potentially form an inter-rod excitation energy transfer (EET) network (Supplementary Fig. 10a). In addition to their close proximity, a high orientation factor (*κ*^2^) has also been shown to indicate increased rates of EET formation between bilin donors and acceptors^25, 40^. Among the outermost PCBs of adjacent rods, we observed that most PCB pairs in rod hexamer 1 (^Rc1^α_2_^81^- ^Ra1^α_2_^81^, ^Rb1^α_1_^81^- ^Rc’1^α_1_^81^ and ^Rb1^α_3_^81^- ^Rc’1^α_3_^81^) had higher *κ*^2^ values than those in hexamer 2, suggesting that the rod-rod EET efficiency might be higher between core-proximal hexamers than between core-distal hexamers (Supplementary Fig. 10b). Moreover, elongated rods may require a longer EET distance to reach the PBS core compared with hemidiscoidal PBS in other species, suggesting that the *G. violaceus* PBS could provide additional EET pathways among rods to compensate for this disadvantage.

Previous studies suggest that energy converged by rods can migrate through bilins in the core before eventually reaching the terminal emitters, α^LCM^ and ApcD^41^. To examine whether this could occur in *G. violaceus,* we constructed EET models from rods to core and within the core based on distances and orientation factors obtained with our cryo-EM structures (Fig. 6a). Different from PBS in crown cyanobacteria and red algae^23, 24, 26^, the *G. violaceus* PBS lacked direct EET pathways from rods to the A/A’ core due to the absence of bottom rods adjacent to the A/A’ basal core cylinder (Supplementary Fig. 11). The estimated EET rate of rod-to-core transfer was lower than that of transfer within the core (Fig. 6a), suggesting a potential bottleneck for overall energy flow across the entire PBS. This scenario aligned well with a previous simulated model of EET in the PBS of *Synechocystis* 6803 ^26^. Specifically, we observed that energy converged from Ra/Ra’, Rc/Rc’, and Rd/Rd’ could be transferred to the C/C’ core, while energy from Rb/Rb’ could be transferred to B/B’ core. Although EET from C/C’ to A/A’ was less efficient than B/B’ to A/A’, a rod-rod EET could allow energy transfer from Ra/Ra’ and Rc/Rc’ to Rb/Rb’ (Fig. 6b), increasing the efficiency of energy transfer to the terminal emitter via B/B’ to A/A’ EET.

**Figure 6.**
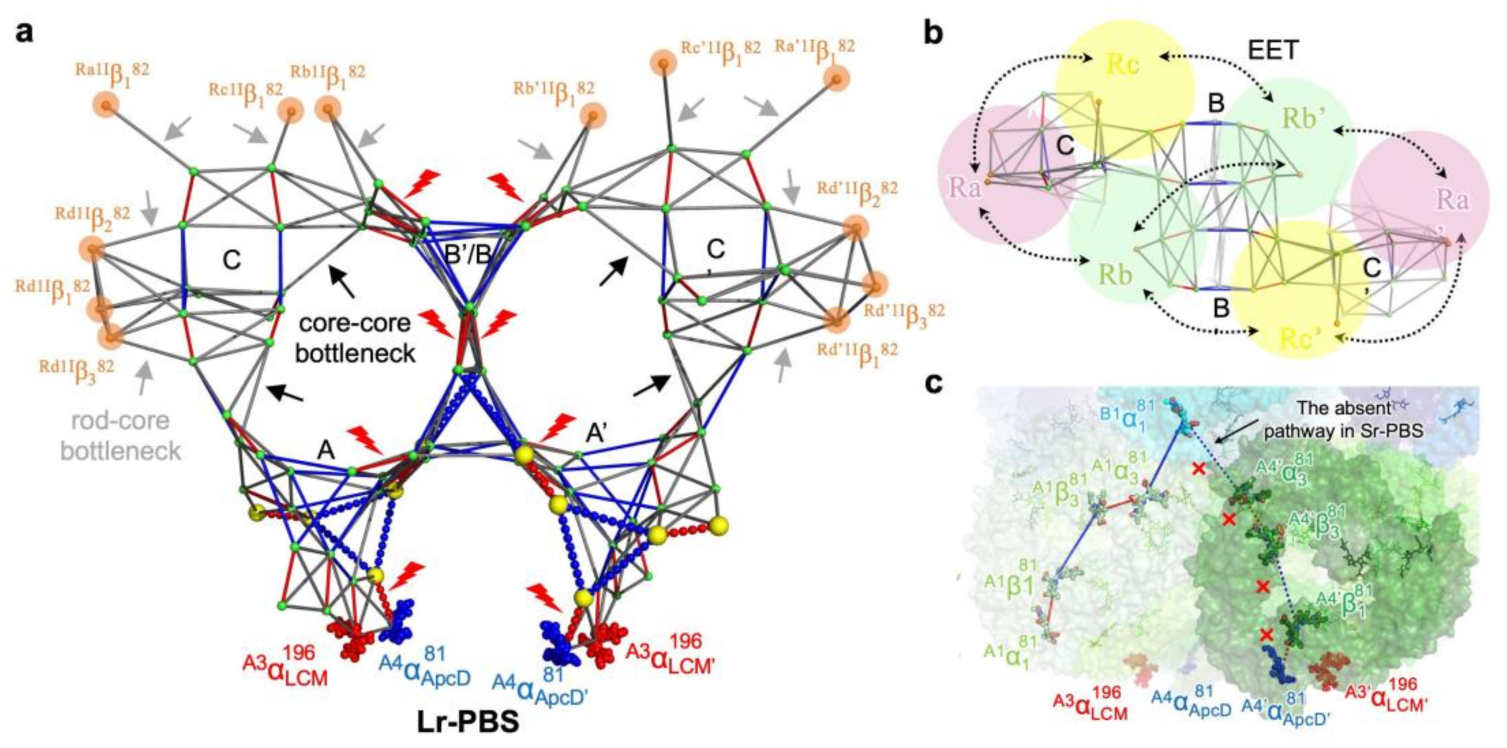
Possible excitation energy transfer (EET) pathways. **a**, Estimated maps of the energy transfer pathways among bilins based on orientation factor (κ^2^) and distances (d). The colors correspond to higher (red), middle (blue) and lower (gray) κ^2^/d^6^ values, which is used to represent the EET rates. The key bilins of rods are shown as sphere in orange color. Bilins that are more than 44 Å are omitted. The gray and black arrows are represented as the rod-core and inner-core bottleneck, respectively. The energy transfer from top B/B’ to bottom A/A’ is high, representing as red lightnings. The absent bilins and pathways in Sr-PBS are shown as yellow sphere and dashes, respectively. The terminal emitters ^A3^α_LCM_^196^ (^A3^α_LCM’_^196^) and ^A4^α_ApcD_^81^ (^A4^α_ApcD’_^81^) are shown in sphere representation. **b**, Top view of the energy transfer pathways. The rods are represented as circles in different colors. EET exchange among the rods is represented by black dash arrows. **c**, The absent of ApcA2-ApcB3-ApcD layer and EET pathways in Sr-PBS is shown as surface representation. The corresponding bilins are shown as sticks.

In addition, compared to the Lr-PBS core, the Sr-PBS core lacked an ApcA_2_-ApcB_3_-ApcD layer, preventing a potential EET pathway through ^B1^α_1_^81^ -^A4’^α_3_^81^ - ^A4’^β_3_^81^ - ^A4’^β_1_^81^- ^A4’^α_ApcD_^81^. These findings suggested that energy from ^B1^α_1_^81^ could only transfer along the A1 layer (^B1^α_1_^81^ - ^A1^α_3_^81^ - ^A1^β_3_^81^ - ^A1^β_1_^81^ - ^A1^α_1_^81^) (Fig. 6c). As the ApcA_2_-ApcB_3_- ApcD layer could not bind to the core due to conformational changes in loop1, loop2, and the bulge of L_CM_ under increased light intensity *G. violaceus* 7421 (Fig. 5), representing a possible adaptation to prolonged exposure to high intensity light conditions, we hypothesized that EET within the core might also correspondingly change to accommodate variations in light intensity.

Chromophore conformation has been shown to change under different light intensities and wavelengths^47^. Here, locally solved APC core structures at ∼3.0 Å and ∼2.6 Å resolution for Lr-PBS and Sr-PBS, respectively, enabled analysis of their PCB conformations (Supplementary Fig. 12a). We observed that most PCBs, except ^A3’^β_3_^81^ and ^A3’^β_1_^81^, exhibited a similar conformation in the conjugated state (Supplementary Fig. 12b and c) wherein the pyrrole rings of ^A3’^β_3_^81^, especially the B and C rings, in Sr-PBS displayed lower coplanarity than those of ^A3’^β_3_^81^ in Lr-PBS. Similarly, the D ring of ^A3’^β_1_^81^ showed lower coplanarity in Sr-PBS compared to that in Lr-PBS, while the coplanarity factors between ring A and ring D of ^A3’^β_1_^81^ and ^A3’^β_3_^81^ were closer to 1 in Lr-PBS than in Sr-PBS. These results suggested that these bilins had lower energy levels in the Lr-PBS core, further suggesting that increased light intensity might induce adaptive changes in bilin energy levels. In addition, although ApcF has been identified as a key subunit determining the efficiency of energy transfer to L_CM_^48, 49^, the *G. violaceus* PBS lacks an ApcF subunit, which is replaced by ApcB^14^ (Fig. 1). However, the conformation and coplanarity factor of the ^A3’^β_1_^81^_ApcB_ in Lr-PBS were comparable to that in ^A2β87^, suggesting a potentially similar function in EET (Supplementary Fig. 12d).

In conclusion, *G. violaceus* 7421 cells grown under low and moderate light intensity exhibit different PBS structures, including an Sr-PBS with short rods lacking an ApcA2-ApcB3-ApcD core layer associated with moderate light intensity, and an Lr-PBS, with long rods and a complete core under low light intensity. These structures suggest that the *G. violaceus* PBS can gradually adapt to high light intensity environments by reducing the length of rods, as well as by altering the structures of loop1, loop2 and the bulge in the first Pfam00427 domain of L_CM_ to prevent ApcA_2_- ApcB_3_-ApcD layer binding to the core. In addition, we solved the structures of linker proteins, L_RC_^91kDa^ and L_RC_^81kDa^, revealing their essential roles in the assembly of bundle-shaped rods. Furthermore, our comparison of these PBS structures in *G. violaceus* 7421 with those of crown cyanobacteria and red algae illustrates an evolutionary trajectory of PBS from thylakoid-free to thylakoid-containing photosynthetic organisms.

## Methods

### Preparation of phycobilisomes

*G. violaceus* 7421 cells were originally provided by Xudong Xu from Institute of Hydrobiology, Chinese Academy of Sciences, Wuhan, Hubei, China. This cyanobacterium might prefer to grow under low light intensity (∼5 μmol photons m^−2^ s^−1^)^50^. *G. violaceus* 7421 cells were grown photoautotrophically at 20°C in BG11 freshwater medium under a fluorescent lamp with a light intensity of 33.8 - 37 μmol photons m^−2^ s^−1^ and supplied with continuously ambient air. Cultures were manually shaken every several weeks. Cells were cultured after 12 months, and then collected for Sr-PBS isolation. For low light growth condition of *G. violaceus* 7421 cells, the cells were grown under the same environments except for the light intensity of ∼5 μmol photons m^−2^ s^−1^. Cells were cultured after 2 months and collected for Lr-PBS isolation. Both Sr-PBS and Lr-PBS were isolated by ultracentrifugation method described early^18^ with little modification. Shortly, cells were broken twice by French Press at 8,000 p.s.i.. Then membranes were incubated with Triton X-100 at a final concentration of 2% for 30 min. Then crude solution was collected immediately after centrifugation for 30 min at 20,000 g. The crude solution was loaded on top of a discontinuous sucrose gradient with slight cross-linking by glutaraldehyde (GA) modified from GraFix method^51^ and spun for 4 h at 120,000 g by using a SW41 rotor on Optima XPN-100 centrifuge (Beckman Coulter). The intact Sr-PBS was on 0.5 M - 0.75 M with blue color. This band was collected and concentrated to a concentration of 1.5 mg/mL. The intact Lr- PBS was on 0.75 M - 1.0 M with purple color. This band was collected under dark and concentrated to a concentration of 0.5 mg/mL. The entire protein isolation process is performed in a dark environment until the proteins are vitrified in the cryo-EM grids.

### Absorption and fluorescence emission spectra

Absorption of the intact PBS was measured between 300–800 nm using an Ultrospec 9000PC ultraviolet–visible spectrophotometer (GE Healthcare). Fluorescence emission spectra were recorded using a UV-800C fluorescence spectrophotometer (HORIBA) at room temperature. After exciting at 450 nm, fluorescence emission was monitored from 500 to 820 nm.

### Cryo-EM sample preparation and data collection

The homemade ultrathin carbon films are preparation by Denton DV-502B (FEI Company). Quantifoil R2/1 (300 meshed) copper grid are covered with the above carbon films for cryo-EM sample preparation, and then glow discharged at low set for 30s by Plasma Cleaner PDC-32G. Then 1.5 μL above phycobilisomes solution with a concentration of 1.5 mg/mL are used for dropping on the copper grid, blot for 3.5s in Vitrobot Mark IV (FEI Company) and then vitrified immediately in liquid ethane cooled by liquid nitrogen.

The cryo-EM data was collected using a 300 kV Titan Kiros Microscopy (FEI) with a K3 Summit direct electron detector (Gatan). A defocus range from -1.5 to -1.8 μm was used. The K3 camera was operated in super-resolution mode, with a calibrated pixel size of 1.0979 Å. Each exposure was 2.56 s and recorded as a movie of 32 frames, with an accumulative dose of 50 electrons per Å^2^ for each stack. GIF was set to a slit width of 20 eV. The data were collected automatically using the AutoEMation2 software^52^. The stacks were motion-corrected by MotionCorr2^53^ with binned twofold.

### Cryo-EM data analysis

The workflow of data analysis for Sr-PBS data is shown in Supplementary Figure 2a. two batches of good Titan cryo-EM micrographs (5,366 and 7,054) were manually screened out. The contrast transfer function (CTF) parameters of each micrograph were estimated by CTFFIND4^54^. Auto-picking by 2D averaged template were performed using Relion3.1.3^55, 56^, which yield 486,556 and 556,066 particles for the next 2D classification, respectively. After several rounds of 2D classification, 227,724 and 509,241 particles were left for the next 3D classification, respectively. After 3D classification, 2 classes and 1 class of the particles with reliable angle distribution were selected for the final reconstruction. At this point, we re-extracted 143,034 and 249,396 particles respectively from the dose-weighted micrographs, and the final resolution of the 3D auto-refinement was 3.97 Å and 3.74 Å, respectively. Then, 2 batches of particles were merged to perform the 3D refinement, CTF refinement and particle polish, and the resolution of the Sr-PBS was 2.98 Å. All particles were transported to CryoSPARC 4.1.1^57^ for further analysis. The overall resolution of the Sr-PBS was 2.80 Å (the center core region can reach to 2.6 Å) after non-uniform refinement and CTF refinement. Since rod regions are blurrier than the center region due to the flexibility, we applied *C*_2_ expansion to the particle dataset, and carried out the particle subtraction and euler angle local search upon 4 local masks for rod a to rod d, which resulted in the improved quality of local maps with resolutions ranging from 2.65 Å and 3.47 Å. The maps for the target regions were extracted from the overall map by Chimera^58^, and the 4 individual masks were created by CryoSPARC 4.1.1. All resolutions were estimated with the gold-standard Fourier shell correlation 0.143 criterion with high-resolution noise substitution.

The workflow of data analysis for Lr-PBS data is shown in Supplementary Figure 2b. Simiar to the process of the Sr-PBS, 2 batch of good Titan cryo-EM micrographs (3,421 and 7,521, respectively) were screened out. The 2D classification, heterogeneous refinement and non-uniform refinement and CTF refinement of these 2 batches of data were performed with CryoSPARC 4.1.1 as illustrated in Supplementary Figure 1b. 188,520 and 417,230 particles are auto-picked by Topaz 0.2.5a^59^ for the next 2D classification. After both 2 rounds 2D classification, 133,795 and 387,011 particles are left for the next heterogeneous refinement, respectively. After both heterogeneous refinement into 2 classes, the class of particles with reliable angle distribution are selected for the final reconstruction. Then we re-extracted 51,049 and 141,765 particles respectively and performed non-uniform refinement, the resolution was 3.78 Å and 3.56 Å, respectively. Then, 2 batches of particles were merged to perform the non-uniform refinement, CTF refinement and local refinement with *C*_2_ symmetry, and the final resolution of the Lr-PBS was 3.37 Å.

### Model building and refinement

We first docked the reported PBS subunits from *G. violaceus* (PDB ID: 2VJR, 2VJT) into the cryo-EM densities using Chimera^58^. Other unknown components of the bundle-shaped PBS were obtained by National Center for Biotechnology Information Nucleotide Database (NCBI) protein blast by homology genes from other cyanobacteria and red algae, including ApcD, L_CM_, L_RC_^91KDa^, L_RC_^81KDa^, L_C_, L_RT_, L_R_^CpcC1^ and L_R_^CpcC2^. The structures of the components were predicted by AlphaFold2^60^. Then, the predicted structures were docked into the cryo-EM maps in Chimera^58^, and modified the main and side chains in Coot^61^ according to the residues with bulky side chain information such as Phe, Tyr, Trp, Arg *e.t.c.* The initial model was completed via iterative rounds of manual building with Coot^61^ and refinement with phenix.real_space_refine^62^. The model was refined with secondary structure and geometry restraints to prevent over-fitting. Notably, the side chains of ApcD are well-fitted with the corresponding map, indicating that the ApcD has been assembled to the PBS core (Supplementary Figure 13). The data collection, model refinement and validation statistics are presented in Supplementary Table 2 and 3. All the figures were prepared in PyMOL (http://pymol.org) or ChimeraX^63^.

### Phylogenetic analysis

The sequences of L_RC_, L_R_ and L_CM_ from thylakoid-lacking cyanobacterium *G. violaceus* 7421 and other crown cyanobacteria and red algae were collected from NCBI (Supplementary Data 1). These full sequences were aligned respectively by Multiple Alignment Mode in ClustalX2^64^, and created by ESPript v.3.0^65^. The phylogenetic relationships of the full sequences of L_RC_s, separated Pfam00427 domains of L_RC_s and L_R_ were analyzed by the maximum likelihood (ML) method in MEGA11 software^66^. To assess the reliability of a phylogenetic tree, 100 bootstrap replicates were resampled. The amino acid substitution model: Jones-Talyor-Thornton (JTT) model was used to simulate the data. Distribution of substitution rates among sites was set as “Uniform Rates”. We used the Nearest-Neighbor-Interchanges (NNI) method to perform a heuristic search of the evolutionary tree. The initial tree for heuristic search were obtained automatically by applying the default Neighbour-Joining (NJ) method. The phylogenetic trees were annotated and visualized in online tool iTOL (https://itol.embl.de/)^67^. The corresponding sequences and tree files have provided in Supplementary Data 1 and 2.

### Estimation of the orientation factor between chromophores

The orientation factor (*κ*) of inter-rods and core was estimated. The *κ* is given by Eq. (1)^25, 41^:

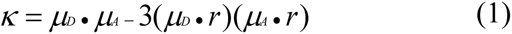

where *μ_D_* and *μ_A_* are the transition dipole moment vectors of the donor (D) and the acceptor (A), respectively. *r* is the distance between D and A. The transition dipole moment vectors of PCBs was determined by referring to ref. ^25, 68^

The Eq. (1) can be estimated by Eq. (2):

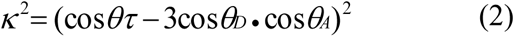

where the 𝜃𝜏 is the angle between *μ_D_* and *μ_A_*, and the 𝜃*_D_* and *θ_A_* are the angels between *μ_D_* and *μ_A_* and the line joining them. The graphic representation can refer to ref. ^25, 41^ Based on the arrangement of the bilins and Eq. (2), *κ*^2^ was calculated using the homemade Python scripts. In brief, firstly, we calculated the distances among adjacent bilins, and the bilins that more than 44 Å are discarded. Second, the 𝜃𝜏, 𝜃*_D_* and *θ_A_* of the remained bilins were calculated and output the *κ*^2^ values in descending order. Finally, the *κ*^2^/*r*^6^ values were calculated to estimated possible energy transfer pathways among these bilins.

### Date Availability

Structure and density map of Sr-PBS is deposited with PDB ID of 9K9W, and map with EMDB ID of EMD-62201. Rod a to Rod d local maps with improved map quality are deposited with EMDB ID of EMD-35801 to EMD-35804, and composite map of bundle-shaped PBS with short rod forEMD-63605. The structure of Lr-PBS is deposited with PDB ID of 9M3J (only containing PBS core), and map of overall PBS with EMDB ID of EMD-63604. Source data are provided with this paper.

## Acknowledgement

We thank Professor X.D. Xu from the Institute of Hydrobiology, Chinese Academy of Sciences for the generous gift of the cyanobacterium, *Gloeobacter violaceus* PCC 7421. We thank Jianlin Lei, Fan Yang, Xiaomin Li and other staff at the Tsinghua University Branch of the National Center for Protein Sciences Beijing for their technical support with cryo-EM and high-performance computing platforms. The authors thank the Protein Chemistry Facility at the Center for Biomedical Analysis of Tsinghua University for sample analysis. We thank Hanzhi Lin and Xijiang Pan for their preliminary research on this project, Wenda Wang for advice on photosynthetic activity analysis, Li Yu and Junmin Pan for advice on regulatory gene expression analysis, Yinchu Wang and Fangqing Zhao for advice on evolutionary analysis. This work was supported by funds from the National Natural Science Foundation of China (32271245 to S.-F.S, 32000848 to J.M. and 32401006 to X.Y.), Young Elite Scientist Sponsorship Program by CAST (2023QNRC001 to X.Y.), China Postdoctoral Science Foundation (2023M741985 to X.Y.). Fundamental Research Funds for the Central Universities of NanKai University (63243138 to J.M.) and Fundamental Research Funds for the State Key Laboratory of the Central Universities of NanKai University (BB042412 to J.M.).

## Author Contributions

S.-F.S. supervised the project; J.M. performed the biochemical and biophysical analysis, cryo-EM sample preparations, cryo-EM data collection and analysis, model building and structure refinement, structural and phylogenetic analysis. X.Y. performed the structural analysis, phylogenetic analysis, chromophore energy transfer calculations; J.M., X.Y. and S.S wrote the initial manuscript; X.Y., S.S. and S.-F.S. analyzed the structures and edited the manuscript.

## Competing interests

The authors declare no competing interests.

## Additional information

**Supplementary information** The online version contains supplementary material. **Correspondence and requests for materials** should be addressed to Jianfei Ma or Sen-Fang Sui.

## Notes

### Competing Interest Statement

The authors have declared no competing interest.

