## Supplemental Information for "Light-induced structural adaptation of the bundle-shaped phycobilisome from thylakoid-lacking cyanobacterium *Gloeobacter violaceus*"

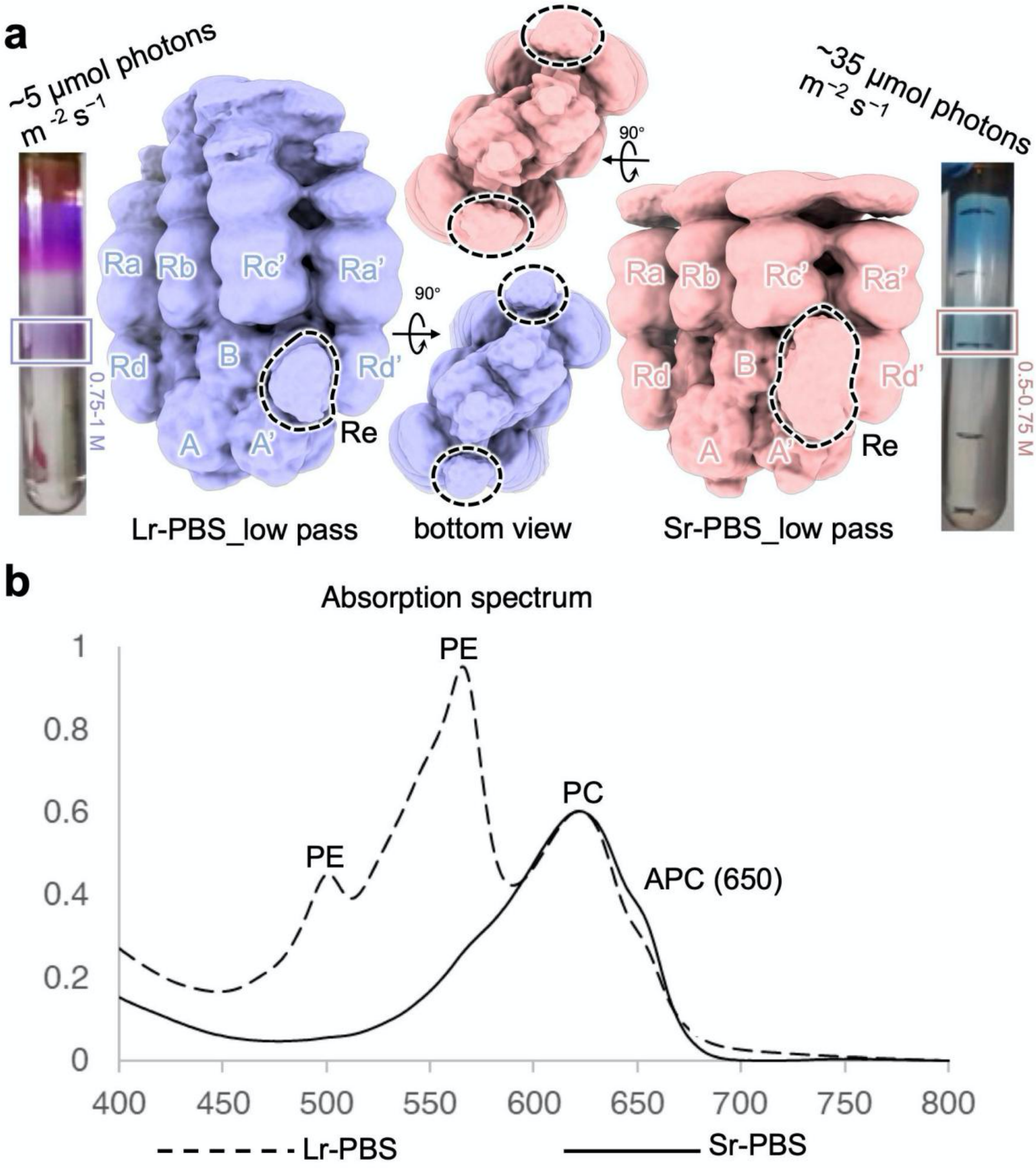


**Supplementary Figure 1 | Preparation, characterization and low-pass filtered structures of the PBSs from *G. violaceus* 7421.**

**a**, Isolation of PBSs using sucrose density gradient centrifugation. The boxes are the sample of the Lr-PBS and Sr-PBS used for single-particle analysis in this study. **b**, Absorption spectrum of the boxes in **a**. The peaks at 500 nm and 560 nm are PEBs. The peaks at 630 nm and 650 nm are PCBs of phycocyanins and PCBs of allophycocyanins, respectively. The dash and solid lines are represented as Lr-PBS and Sr-PBS.

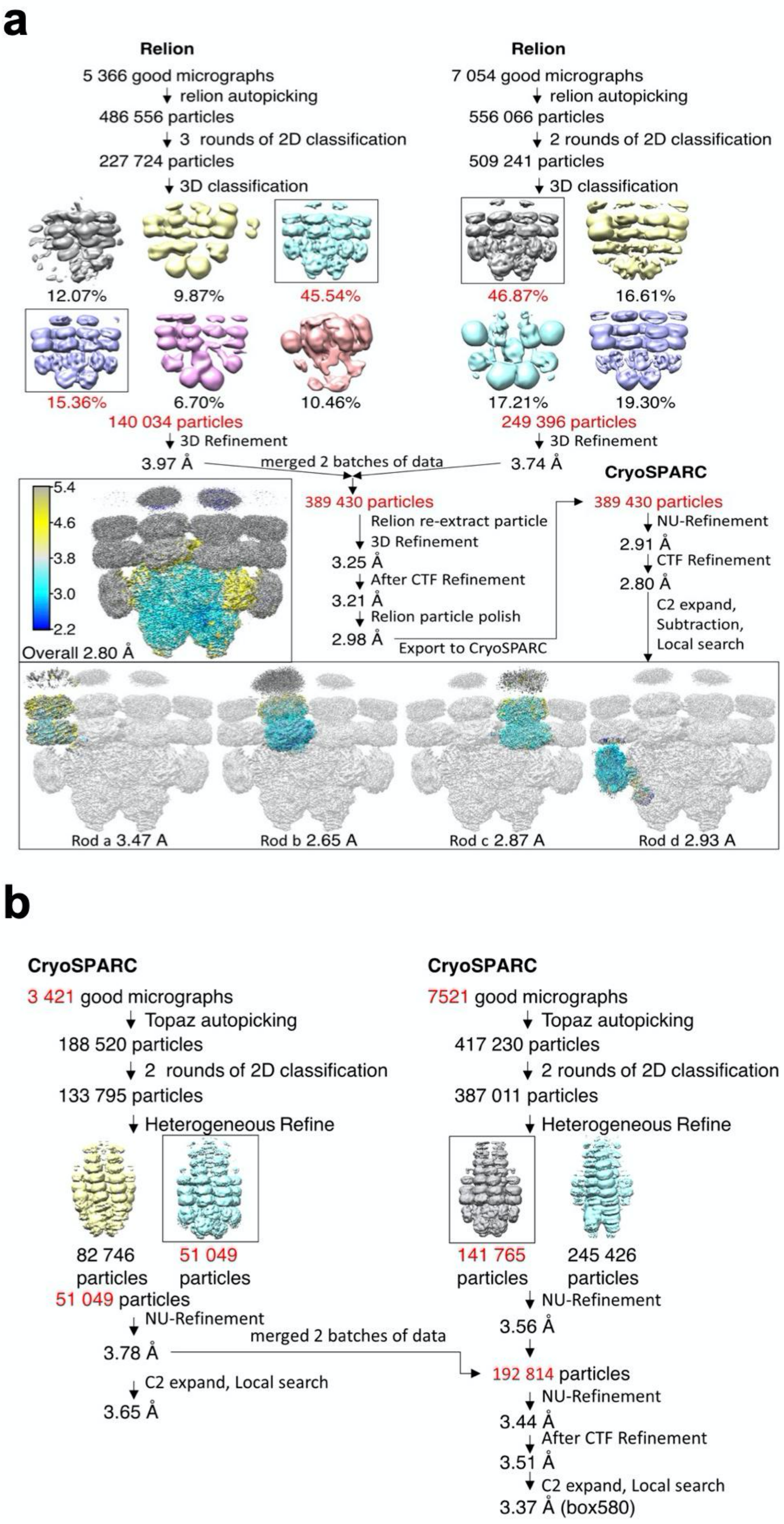
a
Relion
5 366 good micrographs
relion autopicking
486 556 particles
3 rounds of 2D classification
227 724 particles
3D classification
12.07%
9.87%
45.54%
15.36%
6.70%
10.46%
140 034 particles
3D Refinement
3.97 Å
Relion
7 054 good micrographs
relion autopicking
556 066 particles
2 rounds of 2D classification
509 241 particles
3D classification
46.87%
16.61%
17.21%
19.30%
249 396 particles
3D Refinement
3.74 Å
merged 2 batches of data
5.4
4.6
3.8
3.0
2.2
Overall 2.80 Å
389 430 particles
Relion re-extract particle
3D Refinement
3.25 Å
After CTF Refinement
3.21 Å
Relion particle polish
2.98 Å
Export to CryoSPARC
CryoSPARC
389 430 particles
NU-Refinement
2.91 Å
CTF Refinement
2.80 Å
C2 expand,
Subtraction,
Local search
Rod a 3.47 A
Rod b 2.65 A
Rod c 2.87 A
Rod d 2.93 A
b
CryoSPARC
3 421 good micrographs
Topaz autopicking
188 520 particles
2 rounds of 2D classification
133 795 particles
Heterogeneous Refine
82 746 particles
51 049 particles
51 049 particles
NU-Refinement
3.78 Å
C2 expand, Local search
3.65 Å
merged 2 batches of data
CryoSPARC
7521 good micrographs
Topaz autopicking
417 230 particles
2 rounds of 2D classification
387 011 particles
Heterogeneous Refine
141 765 particles
245 426 particles
NU-Refinement
3.56 Å
192 814 particles
NU-Refinement
3.44 Å
After CTF Refinement
3.51 Å
C2 expand, Local search
3.37 Å (box580)

**Supplementary Figure 2 | Data processing of the Sr- and Lr-PBS structures from *G. violaceus* 7421.**

The workflow for the 2D and 3D classifications for cryo-EM data processing. **a**, For Sr-PBS, two parts of data of 5,366 and 7,054 micrographs in two panels are processed by Relion, and reached at 3.97 Å and 3.74 Å resolution individually. Then two parts are merged by Relion and reach 2.98 Å. Finally, data are exported into CryoSPARC to improved map quality at 2.80 Å. The masking strategy for dealing with sub-regions of PBS is enclosed within solid lines. **b**, For Lr-PBS, two parts of data of 3,421 and 7,521 micrographs in two panels are processed by CryoSPARC, and reached at 3.65 Å and 3.56 Å resolution individually. Then two parts are merged, and the final resolution reached to 2.98 Å after NU-Refinement, CTF refinement and local refinement with C2 symmetry. For details, see Cryo-EM data analysis in Methods.

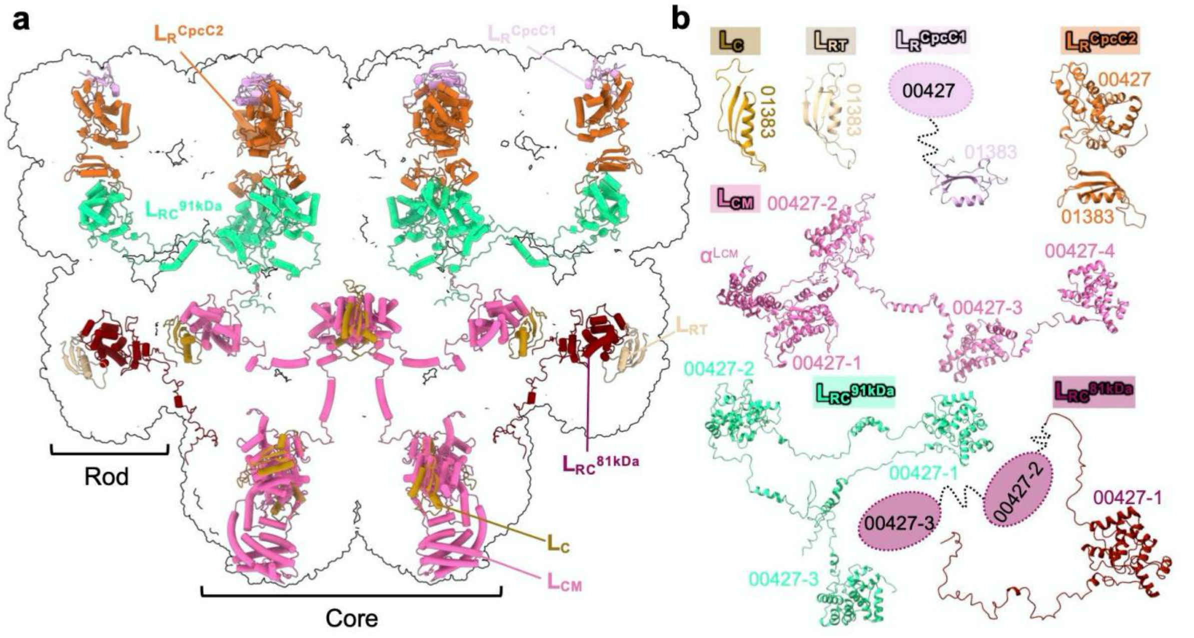


**Supplementary Figure 3 | The distribution of linker proteins.**
**a**, Positions of linker proteins are shown in the PBS. The linkers are shown as cartoon representation in different colors. **b**, Structures of the 7 linkers represented in cartoon.

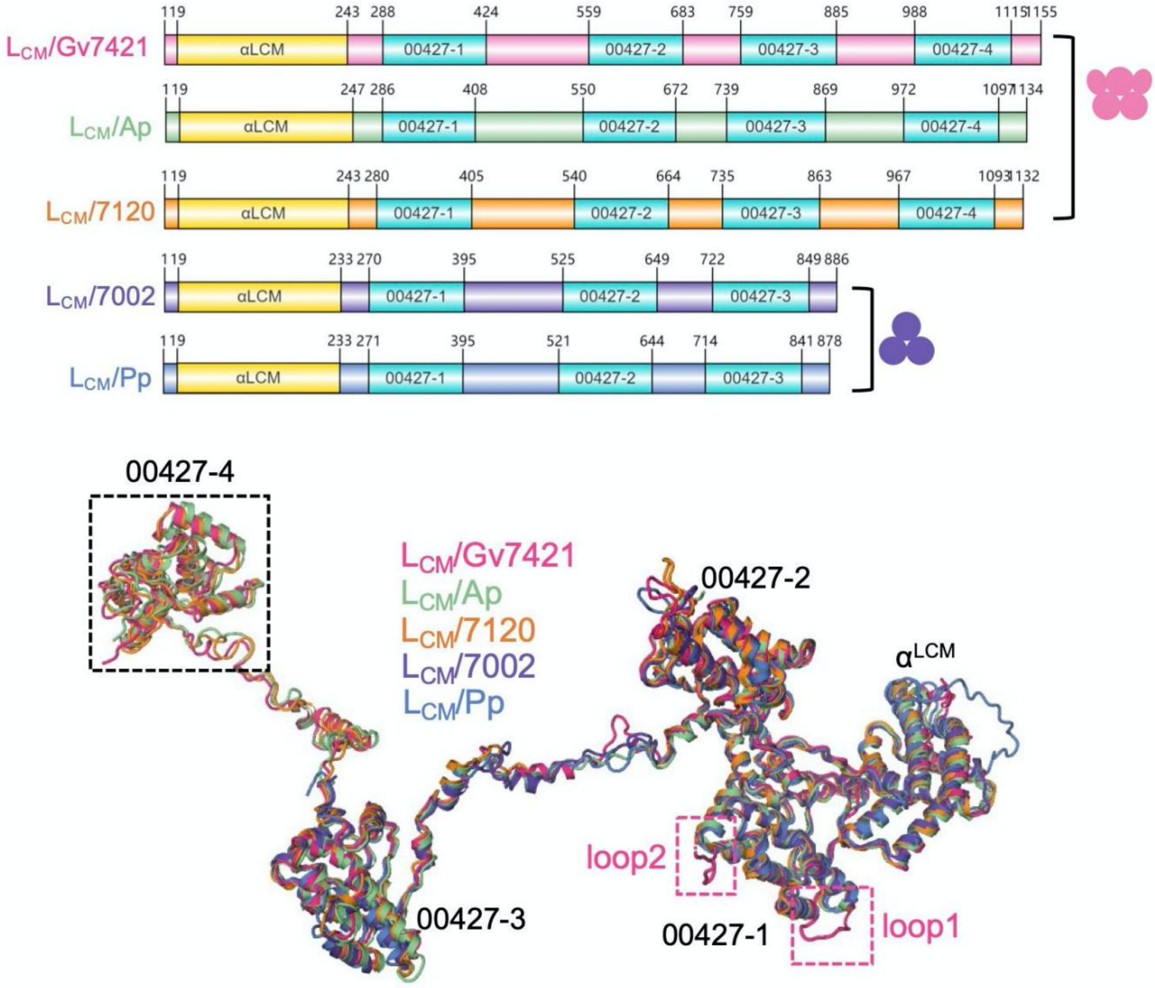


**Supplementary Figure 4 | Structural feature of the $L_{CM}$ from different species.**

**a**, Diagram of the structural elements of $L_{CM}$ from *G. violaceus* 7421, *A. panamensis*, *Nostocaceae* PCC 7120, *Synechococcus* sp. strain PCC 7002 and red algae *P.purpureum*. **b**, Structure of the $L_{CM}$ from *G. violaceus* 7421 (lightmagenta) is superimposed with it from the *A. panamensis* PBS (palegreen), *Nostocaceae* PCC 7120 (orange), *Synechococcus* sp. strain PCC 7002 (purpleblue) and *P.purpureum* (slate). The exclusive loop1 and loop2 of *G. violaceus* $L_{CM}$ are highlighted in the pink dash boxes.

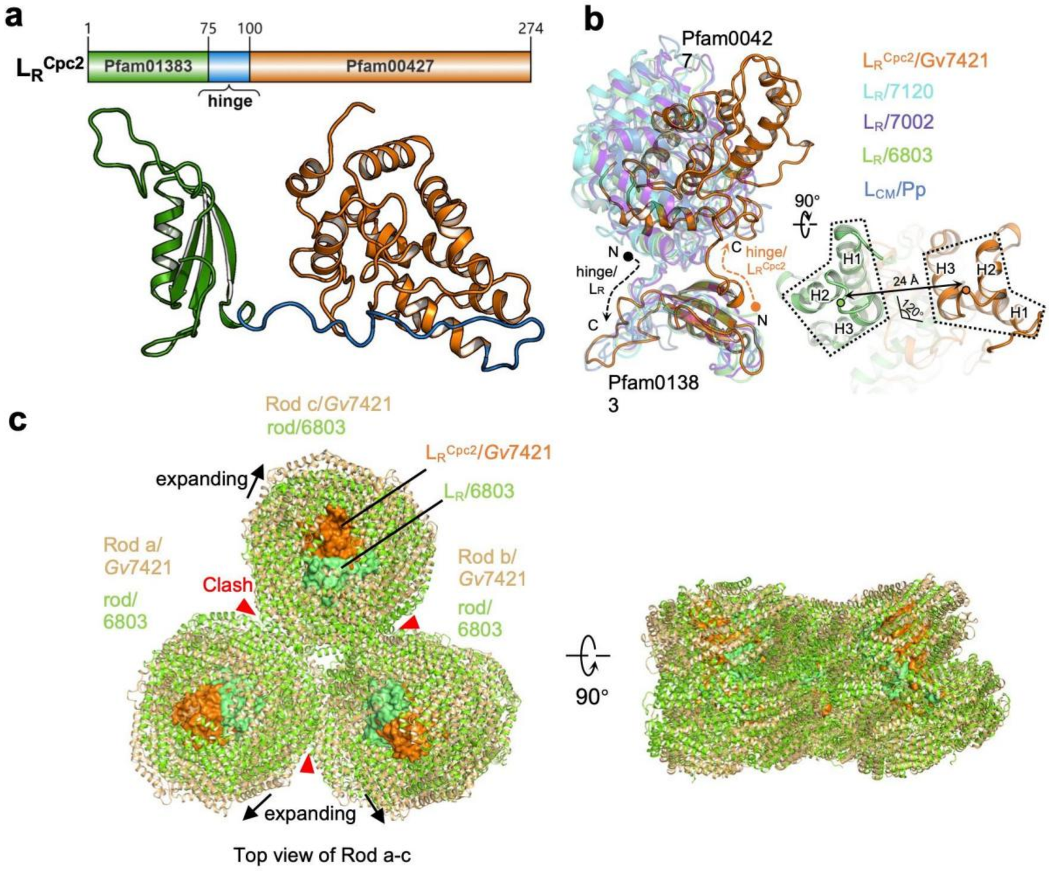


**Supplementary Figure 5 | Structural feature of the $L_R^{CpcC2}$.**

**a**, Top, diagram of the structural elements of the $L_R^{CpcC2}$. Bottom, structure of the $L_R^{CpcC2}$. The N-terminal Pfam01383 domain is represented in forest color. The middle "hinge" is represented in forest color. The C-terminal Pfam00427 domain is represented in orange color. **b**, Structural alignment of the $L_R^{CpcC2}$ from *G. violaceus* 7421 (orange) with the $L_R$ from *Nostocaceae* PCC 7120 (cyan), *Synechococcus* sp. strain PCC 7002 (purpleblue), *Synechocystis* sp. PCC 6803 (limon) and *P.purpureum* (blue based on Pfam01383 domain. The directions of the "hinge" of $L_R^{CpcC2}$ and $L_R$ are from C- to N-terminus and N- to C-terminus, which are labeled as the dash arrow. The Pfam00427 domain of $L_R^{CpcC2}$ is shifted ~24 Å and rotated ~120° compared with the $L_R$. **c**, Structural alignment of a single rod (containing $L_R$) from crown cyanobacterium *Synechocystis* sp. PCC 6803 (limon) and rod a-c (containing $L_R^{CpcC2}$) from *G. violaceus* 7421 (orange). The shift and rotation of Pfam00427 domain of $L_R^{CpcC2}$ could expand the distance among the rods to prevent the clash. The hexamers of rods are shown as cartoon representation. The Pfam00427 domain of $L_R^{CpcC2}$ and $L_R$ is shown as surface representation.

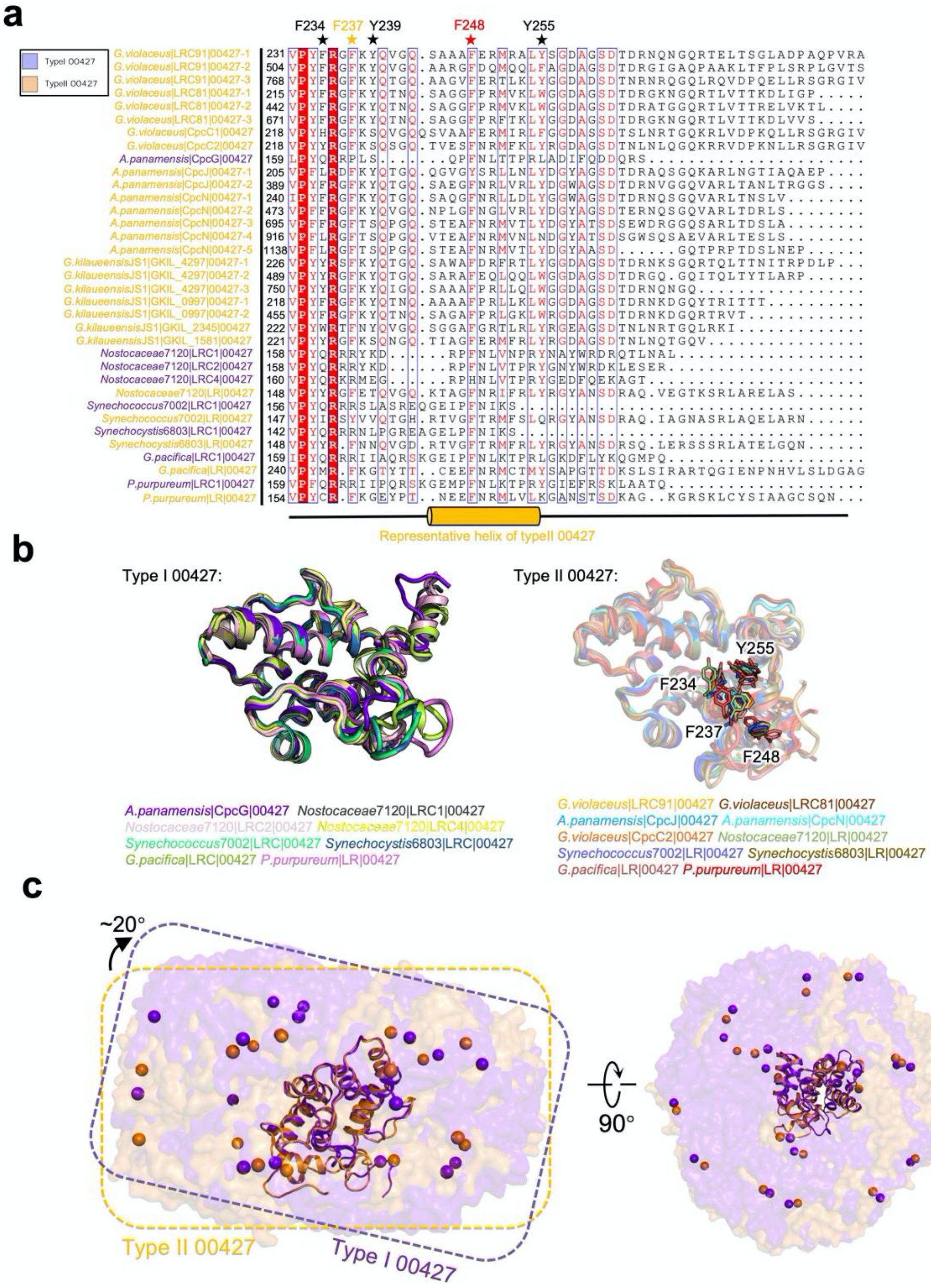

**Supplementary Figure 6 | Sequence and structural alignment of type I and II Pfam00427 domains from different species.**

**a**, Sequence alignment of type I and II Pfam00427 domains of $L_R$ and $L_{RC}$. Type I and II Pfam00427 domains are labeled in purple and light orange, respectively. The key aromatic residues (F234, F237, Y239, F248 and Y255) are marked as star. Among them, the exclusive F237 of type II 00427 domain is coloured in orange, the well conserved residue F248 in both type I and II 00427 domains is coloured in red, the partially conserved residues (F234, Y239 and Y255) are coloured in black. The representative helix of type II 00427 domain is shown as a cylinder in orange. **b**, Left, structural alignment of type I 00427 domain. Right, structural alignment of type II 00427 domain. Key residues are shown as sticks. **c**, Structural alignment of the rod hexamer bound to type I 00427 domain shows ~20° rotation relative to type II 00427. The bilins are shown in sphere.

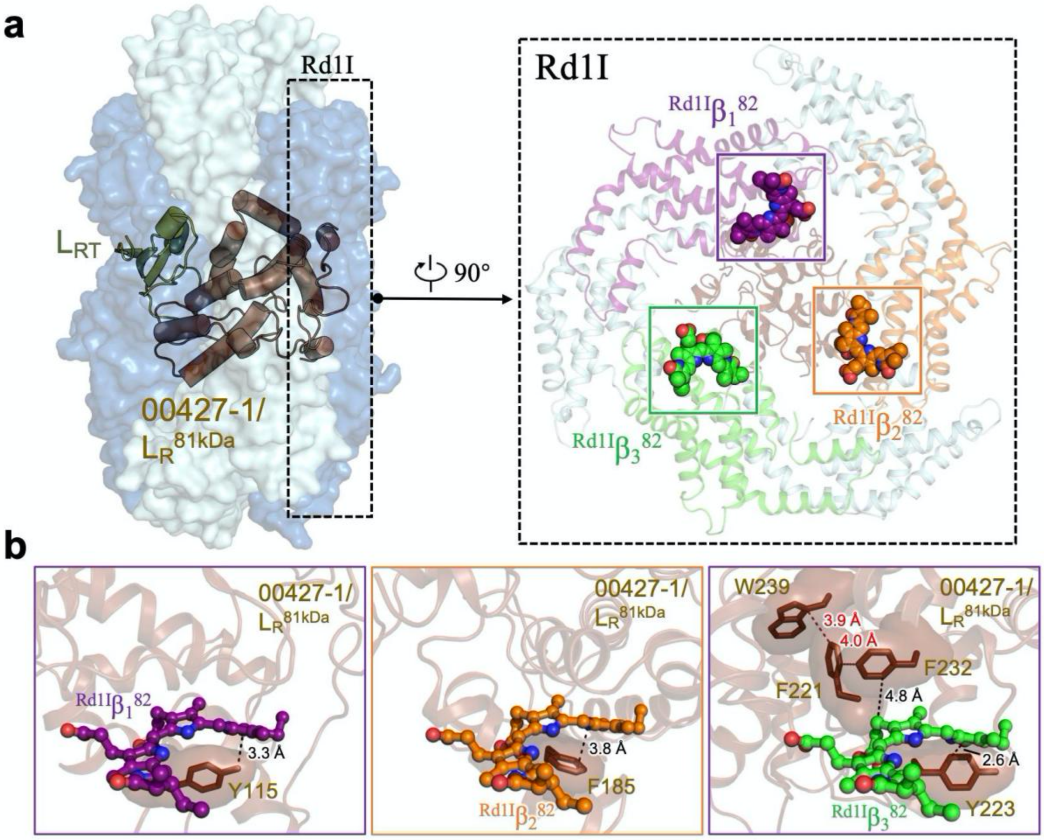


**Supplementary Figure 7 | Interactions of the linker proteins $L_{RC}^{81kDa}$ with chromophores in the rod Rd.**

**a**, Left, overall structure of the rod Rd with the hexamers shown in surface representation and the linker proteins shown in cartoon representation. Right, structure of the layer Rd1I. Proteins and bilins are shown in cartoon and sticks representations, respectively. Three β subunits are coloured differently and the β82 PCBs are boxed and analysed in **b**. **b**, The interactions between the $L_{RC}^{81kDa}$ and the bilins ${}^{Rd1I}\beta_1{}^{82}$, ${}^{Rd1I}\beta_2{}^{82}$ and ${}^{Rd1I}\beta_3{}^{82}$.

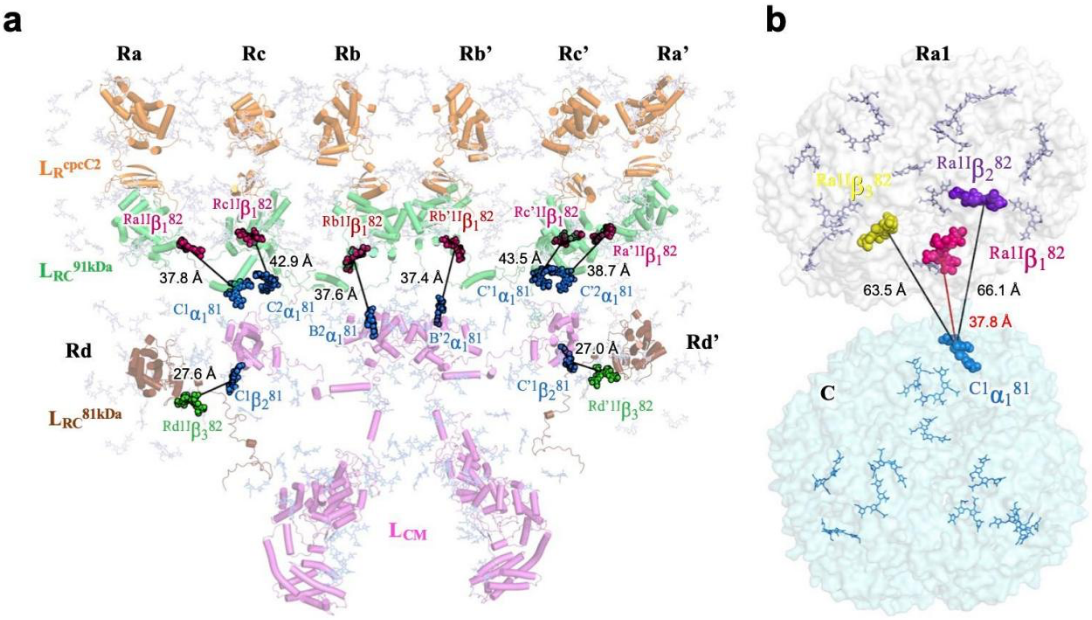


**Supplementary Figure 8 | Distribution of the bilins in rods and core.**

**a**, Positions of the bilins and linker proteins are shown in the PBS. Linkers are shown as cartoon representation in different colors. Bilins are shown in sticks with 70% transparency. The bilins that have the shortest distances between rods and core are highlighted in sphere. The bilins in rods and core are coloured in red, green and blue. The distances are shown as solid lines. **b**, $^{Ra1I}\beta_1^{82}$ has the shortest distance (37.8 Å, red line) between the rod Ra1 and the core compared to $^{Ra1I}\beta_2^{82}$ (66.1 Å, black line) and $^{Ra1I}\beta_3^{82}$ (63.5 Å, black line).

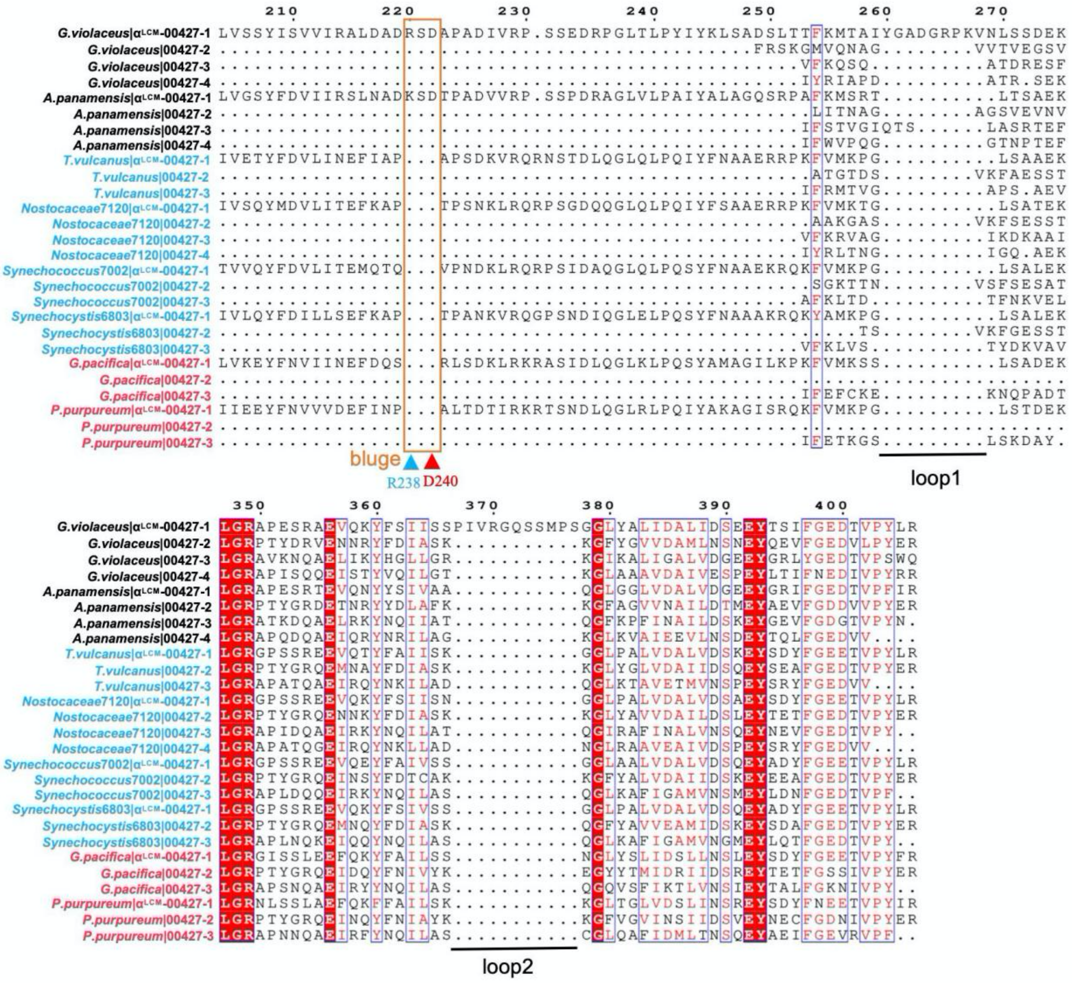

**Supplementary Figure 9 | Sequence alignment of Pfam00427 domains of $L_{CM}$ from different species.**

Sequence alignment of the separated Pfam00427 domains of $L_{CM}$ from thylakoid-lacking cyanobacterium (black font) *G. violaceus* 7421 and *A. panamensis*, and crown cyanobacteria (blue font) *T. vulcanus*, *Nostocaceae* PCC 7120, *Synechococcus* sp. strain PCC 7002, *Synechocystis* sp. PCC 6803 and red algae (red font) *G.pacifica* and *P.purpureum*. The “bulge” is boxed in a orange rectangle. The loop1 and loop2 are highlighted by black lines.

a

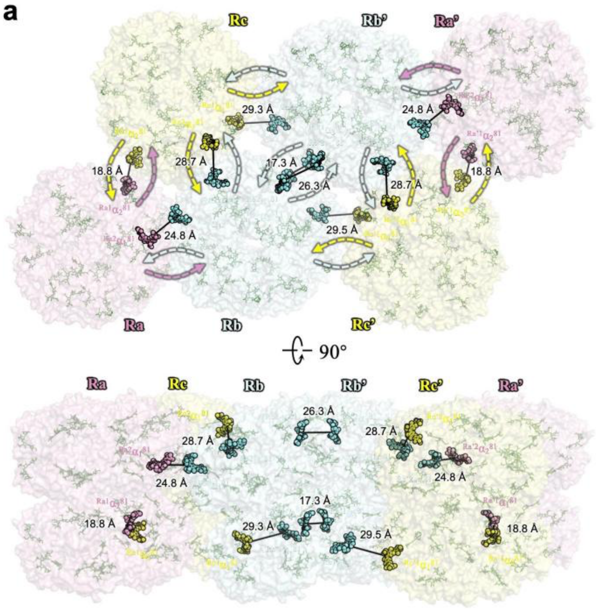


b

| PCB (D/A) | PCB (D/A) | $\kappa^2$ | Distance (Å) |
|---|---|---|---|
| $Rb2\alpha_3^{81}$ | $Rb'2\alpha_3^{81}$ | 0.5480 | 26.3 |
| $Rc1\alpha_2^{81}$ | $Ra1\alpha_2^{81}$ | 0.5363 | 18.8 |
| $Rc'1\alpha_2^{81}$ | $Ra'1\alpha_2^{81}$ | 0.5362 | 18.8 |
| $Rb1\alpha_1^{81}$ | $Rc'1\alpha_1^{81}$ | 0.3044 | 29.5 |
| $Rb'1\alpha_1^{81}$ | $Rc1\alpha_1^{81}$ | 0.3038 | 29.3 |
| $Rb1\alpha_3^{81}$ | $Rb'1\alpha_3^{81}$ | 0.2783 | 17.3 |
| $Rb'2\alpha_2^{81}$ | $Rc'2\alpha_1^{81}$ | 0.0834 | 28.7 |
| $Rb2\alpha_2^{81}$ | $Rc2\alpha_1^{81}$ | 0.0834 | 28.7 |
| $Ra2\alpha_1^{81}$ | $Rb2\alpha_1^{81}$ | 0.0423 | 24.8 |
| $Ra'2\alpha_1^{81}$ | $Rb'2\alpha_1^{81}$ | 0.0423 | 24.8 |

**Supplementary Figure 10 | The inter-rod excitation energy transfer (EET) network.**
**a**, Top, the arrangement of the rods and the inner-bilins in top view. The outmost bilins in adjacent rods that distance within 30 Å are highlighted as sphere representation in different colors. The inter-rod EET is represented as the dash arrows. Bottom, the arrangement of the rods in bottom view. **b**, Estimation of the orientation factor between the highlighted bilins in **a**. The background color of the bilins font corresponds to the rods in **a**. The distance (Å) between a pair of bilins (donor (D) and acceptor (A)), and their orientation factor ($\kappa^2$) is calculated based on Förster theory (See Methods in detail).

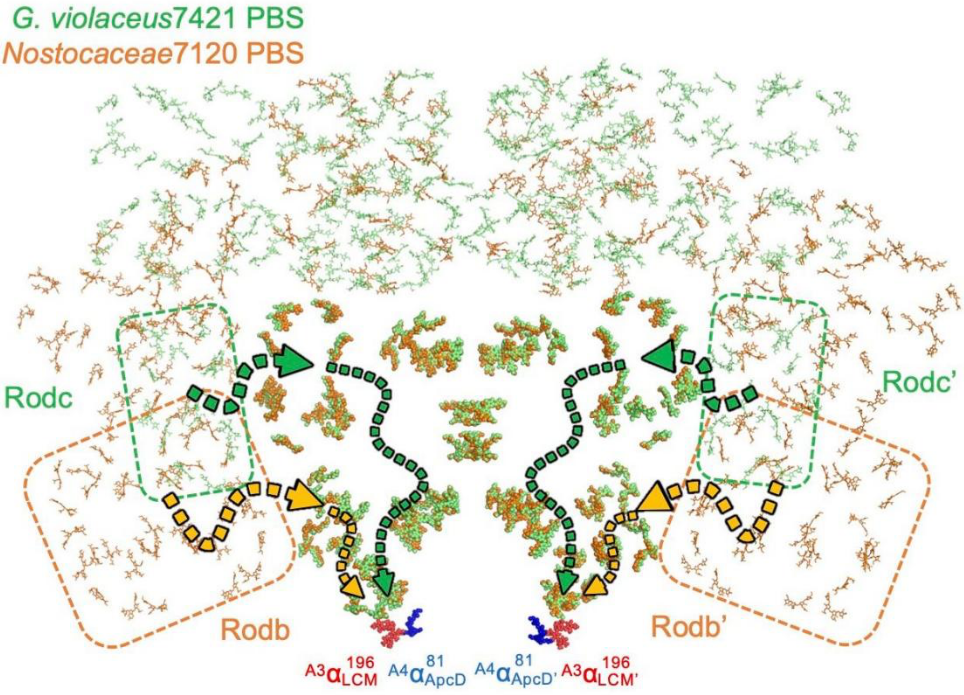


**Supplementary Figure 11 | Arrangement of the bilins from *G. violaceus* 7421 PBS and *Nostocaceae* PCC 7120 PBS.**

Arrangement of the bilins based on the structural alignment of the PBS from *G. violaceus* 7421 (green) and *Nostocaceae* PCC 7120 (orange). The bilins of rods are shown as sticks. The bilins of core are highlighted in sphere representation. Terminal emitters $^{A3}\alpha_{LCM}{}^{196}$ ($^{A3}\alpha_{LCM'}{}^{196}$) and $^{A4}\alpha_{ApcD}{}^{81}$ ($^{A4}\alpha_{ApcD'}{}^{81}$) are coloured in red and blue, respectively. The green and orange dash boxes are represented rod c/c' from *G. violaceus* 7421 PBS and rod b/b' from *Nostocaceae* PCC 7120 PBS, indicating the absence of the bottom rods that neighbored to the basal core A/A' in *G. violaceus* 7421 PBS. In *G. violaceus* 7421 PBS, the energy could migrate via rod c/c' - top cylinders of core - basal core pathways (green arrows), while the energy could transfer from rod b/b' to basal core in *Nostocaceae* PCC 7120 PBS (orange arrows).

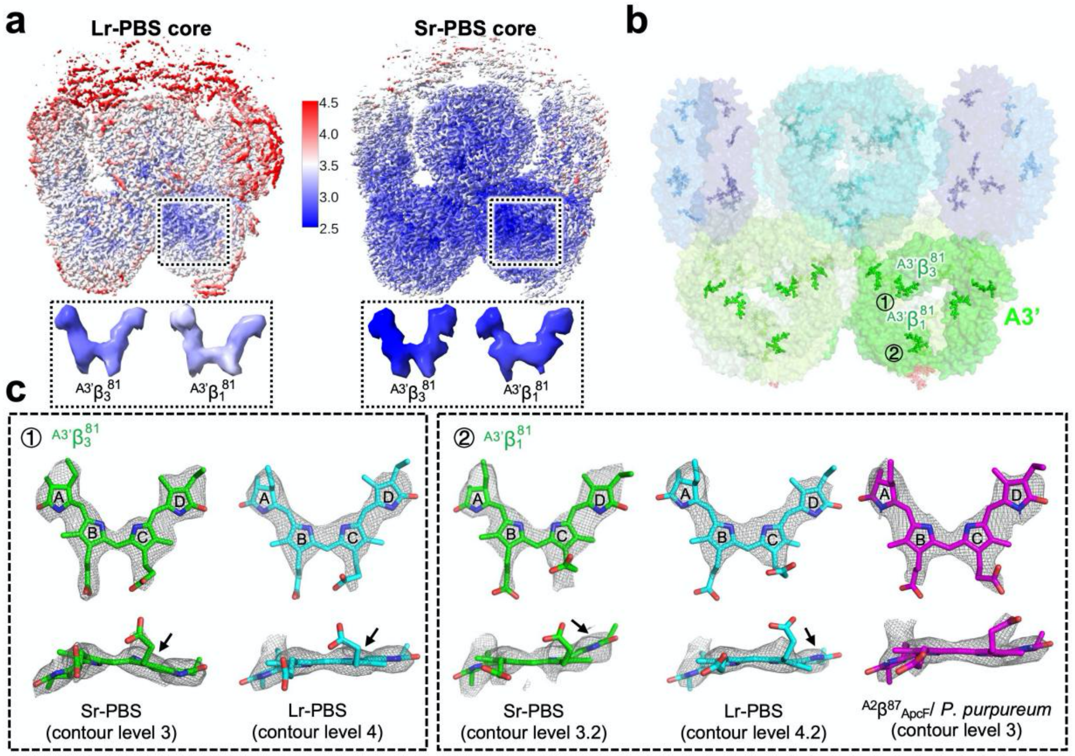


**d**

| | $\Phi_1$ | $\Psi_1$ | $\Phi_2$ | $\Psi_2$ | $\Phi_3$ | $\Psi_3$ | Chemical structure/ Coplanarity statistics |
|---|---|---|---|---|---|---|---|
| $^{A3'}\beta_3{}^{81}$/ Sr-PBS | 28.0 | 128.2 | 27.0 | -0.7 | 157.8 | 22.5 | 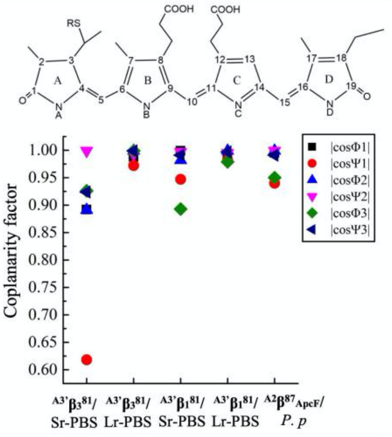 |
| $^{A3'}\beta_3{}^{81}$/ Lr-PBS | -8.9 | 166.5 | 2.6 | 6.6 | 177.2 | -0.8 | |
| $^{A3'}\beta_1{}^{81}$/ Sr-PBS | 1.8 | 161.3 | 10.9 | -4.1 | 153.3 | 7.7 | |
| $^{A3'}\beta_1{}^{81}$/ Lr-PBS | -8.6 | 169.1 | 0.1 | 5.7 | 168.3 | -4.5 | |
| $^{A2}\beta^{87}{}_{ApcF}$/ *P. p* | 5.9 | 160.0 | -0.3 | 0.5 | 161.8 | 7.9 | |



**Supplementary Figure 12 | The densities of the representative bilins show their different coplanarities between Lr- and Sr-PBS**

**a**, Local resolution maps of the Lr-PBS core (left) and Sr-PBS core (right) from *G. violaceus* 7421. The maps were estimated with ResMap and generated in ChimeraX. The local resolutions of the representative bilins are shown in the enlarged boxes. **b**, Structure and bilins arrangement of the PBS core in Sr-PBS. The A3/A3' layers are highlighted in lower transparency (30%). **c**, The densities of the representative bilins in **b** between Lr- and Sr-PBS. The obvious conformation changes of the ring C in $^{A3'}\beta_3{}^{81}$, and ring D in $^{A3'}\beta_1{}^{81}$. The conformation of the $^{A3'}\beta_1{}^{81}$/ApcB in Lr-PBS is similar to $^{A2}\beta^{87}{}_{ApcF}$ from *P. purpureum* PBS. **d**, Dihedral angles of the representative bilins in **c**. The dihedral angles Φ1, Ψ1, Φ2 ... are defined by the plane NA–C(4)–C(5)–C(6), C(4)–C(5)–C(6)–NB, NB–C(9)–C(10)–C(11) ... , etc. |cosΦ1|, |cosΨ1|, |cosΦ2| ... is defined as coplanarity factor.

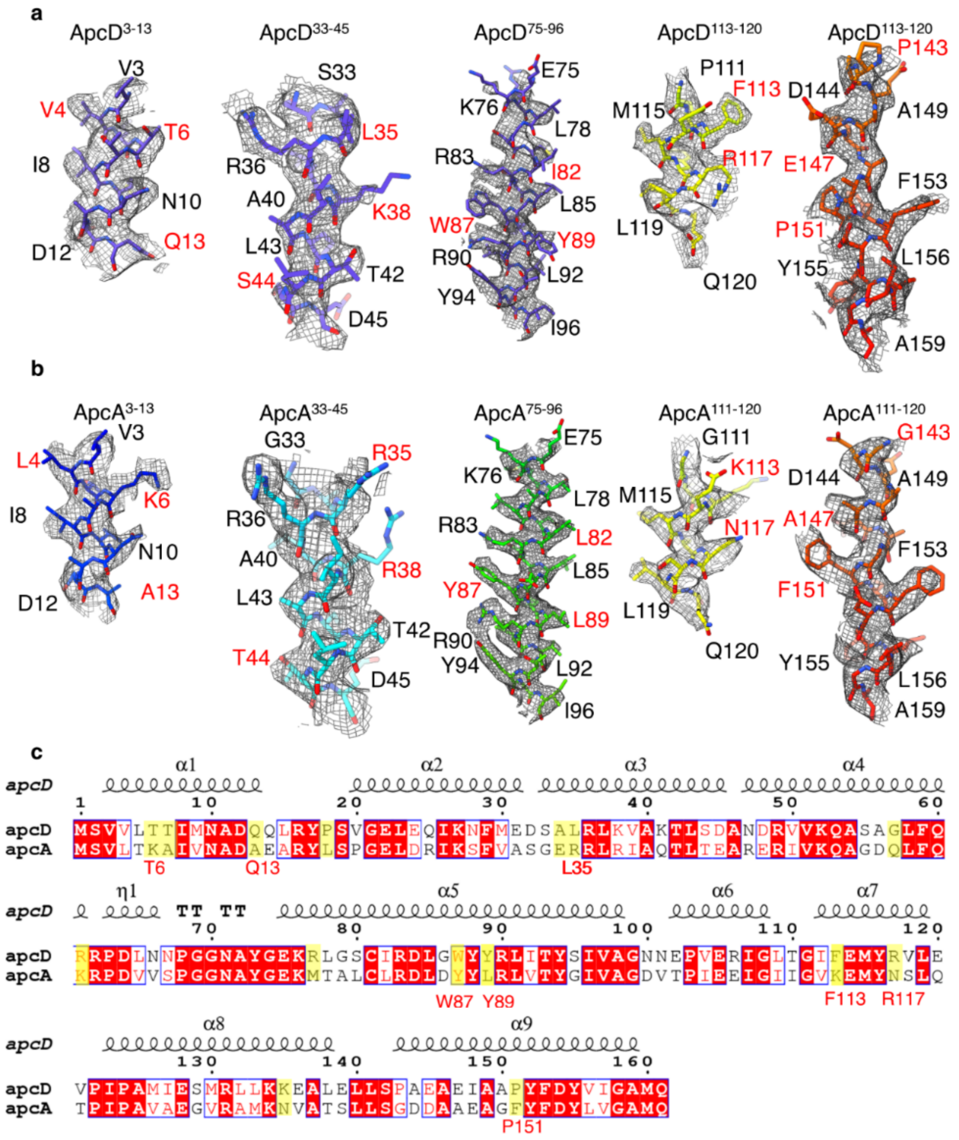


**Supplementary Figure 13| The cryo-EM density maps of ApcD and ApcA, and their refined models.**

a, Selected peptides of ApcD. b, Selected peptides of ApcA. The different side chains are labeled in red color. c, sequence alignment of ApcD and ApcA from *G. violaceus*.

**Supplementary Table 1 |** Summary of proteins and bilins in the modelled structures of Sr-PBS and Lr-PBS. Only 2 layers of $(\alpha^{PC}\ \beta^{PC})_6$ hexamers in Rod a-c/a'-c' are modeled in Sr-PBS. There are no $(\alpha^{PE}\ \beta^{PE})_6$ hexamers (shown in red) resolved at the near atomic-resolution structure.

| | Subunit | Numbers in Sr-PBS structure | Bilin PCB | | Numbers in Lr-PBS core structure | Bilin PCB | |
|---|---|---|---|---|---|---|---|
| | | | Per subunit | Total | | Per subunit | Total |
| 1 | $\alpha^{APC}$ | 40 | 1 | 40 | 44 | 1 | 44 |
| 2 | $\beta^{APC}$ | 42 | 1 | 42 | 48 | 1 | 48 |
| 3 | ApcD | 0 | 1 | 0 | 2 | 1 | 2 |
| 4 | $L_{CM}$_ApcE | 2 | 1 | 2 | 2 | 1 | 2 |
| 5 | $\alpha^{PC}$ | 84 | 1 | 84 | 0 | | |
| 6 | $\beta^{PC}$ | 84 | 2 | 168 | 0 | | |
| 7 | $L_{C}$_ApcC | 4 | | | 6 | | |
| 8 | $L_{RC}$91_Glr1262 | 2 | | | 0 | | |
| 9 | $L_{RC}$81_Glr2806 | 2 | | | 0 | | |
| 10 | $L_{R}$2_CpcC2 | 6 | | | 0 | | |
| 11 | $L_{R}$1_CpcC1 | 6 | | | 0 | | |
| 12 | $L_{RT}$_CpcD2 | 2 | | | 0 | | |
| 13 | $\alpha^{PE}$ | 0 | | | 0 | | |
| 14 | $\beta^{PE}$ | 0 | | | 0 | | |
| | Total | 276 | | 336 | 102 | | 96 |

**Supplementary Table 2 |** Cryo-EM data collection, refinement and validation statistics.

| | #1 Sr-PBS (EMDB-62201) (PDB 9K9W) | #2 Lr-PBS core (EMDB-63604) (PDB 9M3J) |
|---|---|---|
| **Data collection and processing** | | |
| Magnification | 64,000 | 64,000 |
| Voltage (kV) | 300 | 300 |
| Electron exposure (e–/Å$^2$) | 50 | 50 |
| Defocus range (μm) | -1.2 ~ -2.2 | -1.2 ~ -2.2 |
| Pixel size (Å) | 1.091 | 1.091 |
| Symmetry imposed | C2 | C2 |
| Initial particle images (no.) | 1,042,622 | 605,750 |
| Final particle images (no.) | 389,430 | 192,814 |
| Map resolution (Å) | 2.80 | 3.37 |
| FSC threshold | 0.143 | 0.143 |
| Map resolution range (Å) | 2.76~3.49 | 3.37~5.0 |
| **Refinement** | | |
| Initial model used (PDB code) | 2VJR, 2VJT | 2VJR, 2VJT |
| Map sharpening *B* factor (Å$^2$) | -56.2 | -76.6 |
| Model composition | | |
| Non-hydrogen atoms | 382772 | 140502 |
| Protein residues | 48228 | 17848 |
| Ligands | 336 | 96 |
| *B* factors (Å$^2$) | | |
| Protein (mean) | 31.62 | 120.85 |
| Ligand (mean) | 31.86 | 101.38 |
| R.m.s. deviations | | |
| Bond lengths (Å) | 0.021 | 0.010 |
| Bond angles (°) | 2.213 | 1.666 |
| Validation | | |
| MolProbity score | 2.93 | 2.51 |
| Clashscore | 32.62 | 24.91 |
| Rotamer outliers (%) | 6.22 | 2.65 |
| Ramachandran plot | | |
| Favored (%) | 95.28 | 95.66 |
| Allowed (%) | 3.65 | 4.07 |
| Disallowed (%) | 1.07 | 0.27 |

**Supplementary Table 3 |** Summary of model validation for components of the Sr-PBS.

| Molecule* | MolProbity Scores | Ramachandran plot statistics (%) | | | RMS deviations | |
|---|---|---|---|---|---|---|
| | | Favored | Allowed | Outliers | Bonds Length (Å) | Bonds Angles (º) |
| CoreAA′ $L_{CM}$ | 2.72 | 93.07 | 6.25 | 0.65 | 0.014 | 1.567 |
| CoreBC | 2.13 | 96.24 | 3.35 | 0.41 | 0.017 | 1.692 |
| R1$L_{RC}$91 /R1′$L_{RC}$91′ | 2.17 | 95.04 | 4.61 | 0.35 | 0.019 | 1.844 |
| R2/R2′ | 2.01 | 97.14 | 2.83 | 0.02 | 0.019 | 1.702 |
| R3/R3′ | 2.73 | 96.47 | 3.37 | 0.16 | 0.024 | 2.146 |
| R4$L_{RC}$81 /R4′$L_{RC}$81′ | 2.37 | 95.27 | 4.39 | 0.34 | 0.014 | 1. 932 |

* CoreAA′$L_{CM}$ contains all α-subunits, β-subunits in core A, core A′, and $L_C$1/$L_C$1′, $L_{CM}$/$L_{CM}$′; CoreBC contains all α-subunits, β-subunits in core B, core C, core C′, and $L_C$1/$L_C$1′; each rod (R1$L_{RC}$91/R1′$L_{RC}$91′ through R4$L_{RC}$81/R4′$L_{RC}$81′) contains all α-subunits, β-subunits and linker proteins in the rod; R1$L_{RC}$91/R1′$L_{RC}$91′ contains α-subunits, β-subunits, and $L_{RC}$91/$L_{RC}$91′, $L_R$CpcC2/ $L_R$CpcC2′ and $L_R$CpcC1/$L_R$CpcC1′; R2/R2′ contains α-subunits, β-subunits, and $L_R$CpcC2/ $L_R$CpcC2′ and $L_R$CpcC1/$L_R$CpcC1′; R3/R3′ contains α-subunits, β-subunits, and $L_R$CpcC2/ $L_R$CpcC2′ and $L_R$CpcC1/$L_R$CpcC1′; R4$L_{RC}$81/R4′$L_{RC}$81′ contains α-subunits, β-subunits, and $L_{RC}$81/$L_{RC}$81′, $L_{RT}$CpcD2/ $L_{RT}$CpcD2′.
